# miR-124 regulates *Notch* and *NeuroD1* and to mediate transition states of neuronal development

**DOI:** 10.1101/2021.12.10.471989

**Authors:** Kalin D. Konrad, Jia L. Song

## Abstract

MicroRNAs (miRNAs) regulate gene expression by destabilizing target mRNA and/or inhibiting translation in animal cells. The ability to mechanistically dissect the function of miR-124 during specification, differentiation, and maturation of neurons during development within a single system has not been accomplished. Using the sea urchin (*Strongylocentrotus purpuratus*) embryo, we take advantage of the manipulability of the embryo and its well-documented gene regulatory networks (GRNs). We incorporated NeuroD1 as part of the sea urchin neuronal GRN and determined that miR-124 inhibition resulted in decreased gut contractions, swimming velocity, and neuronal development. We further integrated post-transcriptional regulation of miR-124 into the neuronal GRN. Inhibition of miR-124 resulted in increased number of cells expressing transcription factors associated with progenitor neurons and a concurrent decrease of mature and functional neurons. Results revealed that miR-124 regulates undefined factors early in neurogenesis during neuronal specification and differentiation in the early blastula and gastrula stages. In the late gastrula and larval stages, miR-124 regulates *Notch* and *NeuroD1*. Specifically, miR-124 regulates the transition between neuronal differentiation and maturation, by directly suppressing *NeuroD1*. Removal of miR-124’s suppression of *NeuroD1* results in increased mature neurons with decreased Synaptagmin B-positive mature, functional neurons. By removing both miR-124 suppression of *NeuroD1* and *Notch*, we were able to phenocopy miR-124 inhibitor induced defects. Overall, we have improved the neuronal GRN and identified miR-124 to play a prolific role in regulating various transitions of neuronal development.

## Introduction

The sea urchin larva has three neuronal centers: the apical organ with ganglionic organization analogous to the vertebrate central nervous system; the ciliary band that coordinates larval swimming, analogous to the peripheral nervous system; and gut neurons that mediate contraction of the digestive system (Fig. 1A). Although the body plan and neuronal organization of deuterostomes are diverse, developmental mechanisms that mediate the specification and differentiation of metazoan nervous systems share striking similarities at the molecular level. It has been observed that sea urchin neuronal-specific *Pou4f2* (*Brn*) can functionally replace *Pou4f2* in mice, revealing a strong level of conservation in neuronal development across the species^1^. Both vertebrate and sea urchin embryos use the FGF signaling pathway to initiate neurogenesis^2–5^, Nodal and BMP pathways to restrict dorsal-ventral neuronal regions^6–9^, Wnt signaling to suppress neuronal development^10–13^, and the Delta/Notch pathway to mediate classical lateral inhibition that result in Delta-expressing differentiated neurons^14–16^. Additionally, Sox transcription factors, Pou/Brn, and Elav are all conserved proteins used in the specification, differentiation, and maturation of neurons^1, 17–20^. Thus, the sea urchin embryo uses evolutionarily conserved transcription factors (TFs) and signaling pathways to set up the nervous system.

**Figure 1.**
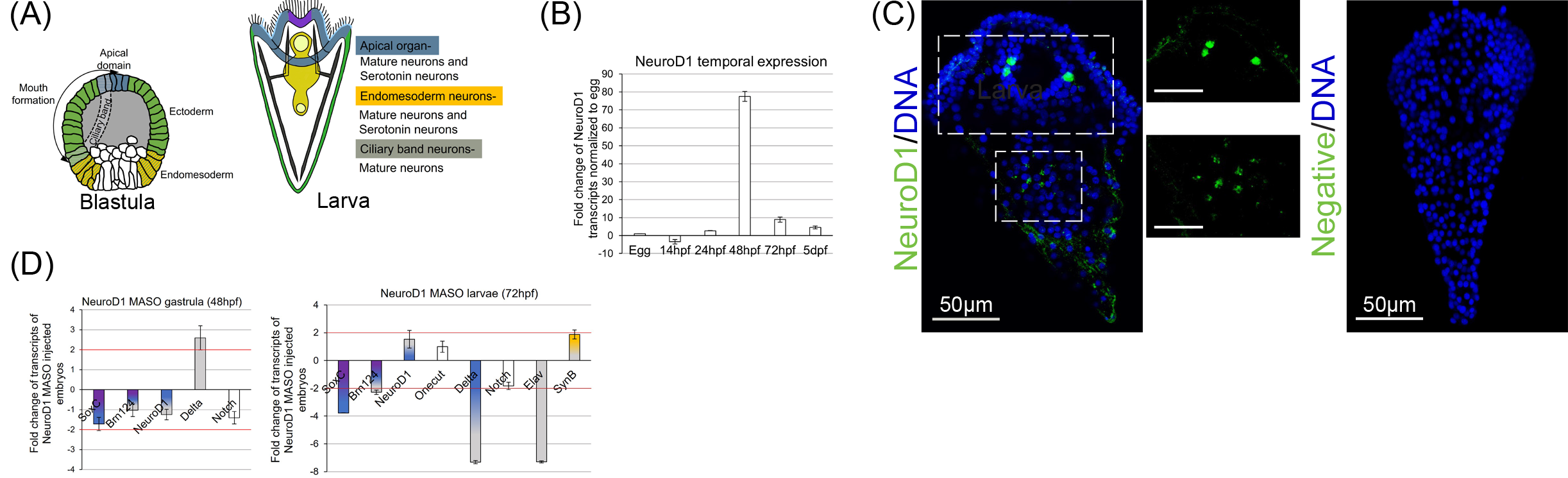
NeuroD1 regulates genes involved in the neuronal GRN. (A) Three neuronal domains are specified during the blastula stage. Neuronal progenitors will differentiate and mature into functional neurons in the larvae stage. (B) 200 embryos were collected during the major developmental time points (Egg (0hpf), morula (14hpf), blastula (24hpf), gastrula (48hpf), larval (72hpf), and 5 days post fertilization (dpf)). 2 embryo equivalents were tested using qPCR. *NeuroD1* expression was normalized to ubiquitin and then to the egg. NeuroD1 expression is highest during the gastrula stage. SEM error bars are graphed. (C) Sea urchin larvae were immunolabled with NeuroD1 (green) and counterstained with DAPI (blue). NeuroD1 is localized to presumptive ganglia, neurofilaments and perinuclearly in cells of the hindgut. (D) 100 of control MASO or NeuroD1 MASO injected embryos were collected for qPCR at gastrula and larval stages. Transcripts of genes involved in neuronal development are examined. NeuroD1 MASO resulted in at least 2-fold increase of *Delta* at the gastrula stage and at least 2-fold decrease of *SoxC*, *Brn1/2/4*, *Delta*, and *Elav* at the larval stage. 3-5 replicates. SEM error bars are graphed.

The neuronal progenitor specification in the sea urchin apical organ (serotonin expressing neurons) coincides with *Foxq2* and *SoxC* expression^21, 22^. In the apical domain early in development, SoxB1 will activate *Foxq2* and *SoxC*^23, 24^. Foxq2 will inhibit canonical Wnt signaling (Wnt6) that will specify the anterior neuroectoderm^24, 25^. Later, the *SoxC*-positive neuronal progenitors also express *Brn1/2/4*^22^. Once the neuronal *SoxC*- positive progenitors undergo their last mitotic division mediated by Delta/Notch signaling, the differentiated neuron becomes *Brn1/2/4* and *Delta* positive^22^. Differentiated, mature neurons in the ciliary band and apical organ express *Elav*^22^, which is an RNA binding protein important to stabilize transcripts that regulate axonal targeting and synaptic growth^19, 20^. The mature and functional neuron will express Synaptotagmin B (SynB), which is part of the SNARE family that mediates neurotransmitter release of synaptic vesicles^26^. A subpopulation of those mature neurons in the apical neuroectoderm also express serotonin^27, 28^. Foxq2 is an important early TF for specifying the apical domain and its restricted expression early in development is critical in proper development of serotonergic neurons^29, 30^. Serotonin is a neurotransmitter important for mediating larval gut contractions, early swimming, and feeding behavior^31, 32^.

The ciliary band consists of thicken layers of monociliated epithelial cells where most of the neurons reside^33, 34^. These layers of cells within the ciliary band are important for proper neuronal connection and activity^28, 35^. The ciliary band is formed when a ventral-dorsal boundary is set up during the blastula stage by Nodal and BMP2/4 signaling pathways^29, 33^. The boundary where these signaling pathways are inactive will allow for the expression of *Onecut (Hnf6)*, which is expressed by specialized cells that makes up the ciliary band where neuronal connections are formed^33, 36–39^.

The third domain of neurons resides in the tripartite gut to mediate muscular contractions for feeding^40^. Each compartment of the tripartite gut is separated by mesodermally-derived sphincters: the cardiac sphincter separates the foregut and the midgut; and the phyloric sphincter separates the midgut and the hindgut^41^. The neurons that reside in the gut and pharynx are endodermally-derived^40^. Less is known about the enteric neurons; however, key transcription factors, *SoxB1*, *Delta*, and *SoxC*, are expressed in the endomesoderm during the blastula stage^24^. *SoxB1, Six3,* and *Nkx2-3* expression in the endomesoderm could specify the foregut as neuroenderm but this had not been proven^40^. Recently, it has been shown that the opening of the pyloric sphincter is responsive to light, as a result of released serotonin that bind to the receptors in the midgut to mediate contraction^42^. During the larval stage, in response to calcium influx and release of different neurotransmitters, neurons in these three neuronal domains mediate swimming and feeding behavior which is vital for proper development.

NeuroD (NeuroD1, NeuroD2, and NeuroD6) TFs are a member of the neuronal lineage basic helix-loop-helix (bHLH) family that regulate the transition from neuronal differentiation to maturation in vertebrate systems, cell lines, *Drosophila melanogaster*, and *Caenorhabditis elegans*^43–53^. However, none of the NeuroD members has been incorporated into the sea urchin neuronal GRN. NeuroD1, specifically, has been demonstrated to be expressed early in mammalian development to regulate neuronal development, suggesting that it is a good candidate to be incorporated into the sea urchin neuronal GRN^48, 50, 51, 53^.

In several organisms, miR-124 is expressed in neuronal tissues and plays an evolutionarily conserved function in regulating the balance between proliferation and differentiation of the nervous system during development^52, 54–59^. Although the function of miR-124 has been examined in several animals^56, 60, 61^, a systematic and comprehensive understanding of miR-124’s role in neuronal specification, differentiation, and maturation in a single animal is still lacking. The sea urchin embryo serves as a powerful model to integrate post-transcriptional regulation of neurogenesis because this embryo can be experimentally manipulated to follow neurogenesis throughout development^6, 22^. The sea urchin embryo contains ∼50 miRNAs compared to the ∼500 of miRNAs identified in humans and mice^62–65^. The low complexity of miRNAs, with a single miR-124, compared to the three different copies in the mouse, makes the sea urchin embryo a tractable model to examine the function of miR-124^66–69^.

In this study, functional studies of *NeuroD1* indicate that NeuroD1 regulates *Brn1/2/4*, *SoxC*, *Delta*, and *Elav*. We determined that miR-124 inhibition resulted in decreased gut contractions and swimming velocity, potentially due to its indirect regulation of serotonin levels. In addition, inhibition of miR-124 resulted in increased number of cells expressing TFs associated with progenitor neurons, such as *FoxQ2, SoxC*, and *Brn1/2/4*, and a concomitant decrease of serotonin-expressing neurons and SynB positive functional neurons. We found that miR-124 regulates undefined factors early in neuronal specification and differentiation during the early blastula and gastrula stages, suppresses *Notch* to mediate the last mitotic division leading to functional neurons, and suppresses *NeuroD1* to mediate neuronal transition from differentiation to maturation in late gastrula and larval stages. Furthermore, we find that miR-124’s suppression of *NeuroD1* and *Notch* is sufficient to phenocopy miR-124 inhibitor-induced defects, indicating that miR-124 fine-tunes these factors to control neuronal development. Overall, we were able to integrate NeuroD1 into the sea urchin neuronal GRN and systematically define miR-124’s regulatory role throughout neurogenesis by identifying its targets within the neuronal GRN.

## Results

### NeuroD1 regulates *SoxC*, *Brn1/2/4*, *Delta* and *Elav*

NeuroD1 has been shown in other systems to be an important TF that mediates proper neuronal development, by promoting neuronal differentiation in progenitor cells, as well as in reprogramming differentiated non-neuronal cells into neurons^46, 48, 70^. We bioinformatically identified *NeuroD1* transcript to contain one miR-124 binding site. Thus, prior to examining miR-124’s post-transcriptional regulation of neurogenesis, we first examined the function of NeuroD1, which has an evolutionarily conserved role in neurogenesis in other organisms^20, 49–51, 53, 71^. By performing quantitative polymerase chain reaction (qPCR), we determined that the expression of *NeuroD1* is low in early developing embryos and peaks at gastrulation, followed by decreased expression during the larval stage (Fig. 1B). Protein alignment analysis indicates that the sea urchin and human NeuroD1 proteins share overall 33.7% identity and 85.1% identity within the DNA binding domain region (Fig. S1). Using a NeuroD1 antibody developed against the human NeuroD1, we determined that this antibody cross-reacted with the sea urchin NeuroD1 (Fig. S2A, S2B). The sea urchin NeuroD1 protein is expressed in presumptive ganglia and neurofilament structures in the ciliary band and perinuclearly in the cells of the gut (Fig. 1C).

To test the function of NeuroD1, we performed loss-of-function study of *NeuroD1* with a translational morpholino (NeuroD1 MASO). NeuroD1 protein was significantly reduced in NeuroD1 MASO injected embryos compared to the control MASO injected embryos, indicating efficacy of the NeuroD1 knockdown and specificity of the NeuroD1 antibody (Fig. S2D). To determine the role of NeuroD1 during neurogenesis, we assayed transcript levels of key TFs and genes within the neuronal GRN in NeuroD1 MASO and control MASO injected embryos at the time when *NeuroD1* is expressed in gastrula and larval stages (Fig. 1D). We observed that during the gastrula stage, the expression of *Delta* was increased 2-fold (Fig. 1D). During the larval stage, the expression levels of *SoxC, Brn1/2/4, Delta,* and *Elav* were decreased at least 2-fold (Fig. 1D). These results indicate that NeuroD1 represses *Delta* in the gastrula stage and activates *SoxC*, *Brn1/2/4*, *Delta,* and *Elav* in the larval stage. Based on these observations, we place NeuroD1 upstream of *SoxC*, *Brn1/2/4*, *Delta* in the neuronal GRN and *Elav* in late gastrula to larval stages.

### miR-124 is enriched in the ciliary band where neurons reside

With NeuroD1 integrated to the neuronal GRN, we then investigate the expression pattern of miR-124 throughout development. We collected and fixed eggs and embryos at different developmental time points and examined the expression of miR-124 by using fluorescent RNA *in situ* hybridization (FISH). Results indicate that miR-124 is not detectable until the morula stage (Fig. 2A). In the blastula and gastrula stages, miR-124 is expressed ubiquitously. Later in the larval stage, miR-124 is enriched in the ciliary band (Fig. 2A). Specifically, miR-124 is expressed in basal epithelial cells juxtaposed to epithelial cells that express *Onecut* (*Hnf6),* where mature neurons (SynB-positive neurons) reside (Fig. 2B)^28, 36, 72^.

**Figure 2.**
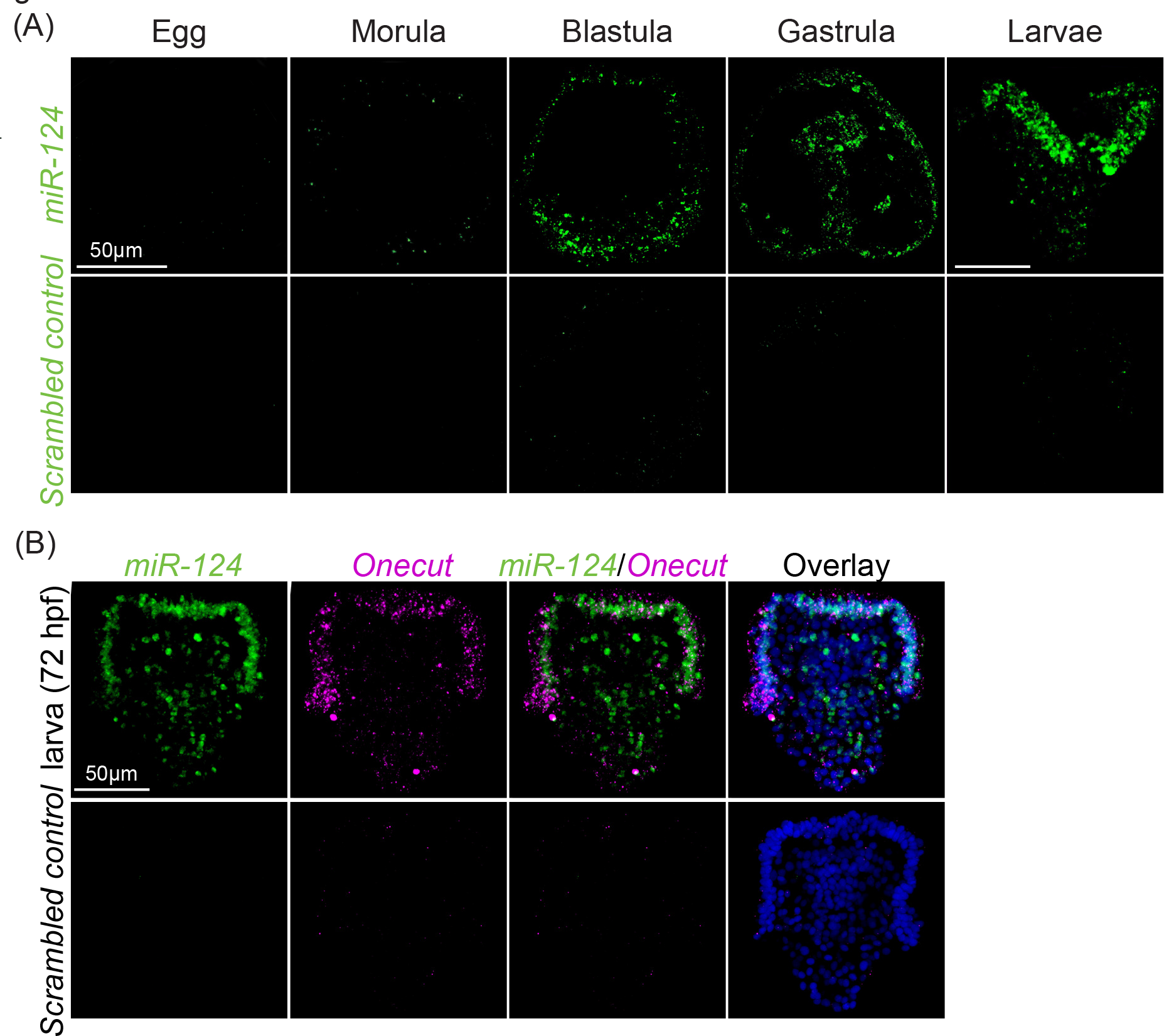
miR-124 is enriched in ciliary band. (A) Embryos were hybridized with miR- 124 LNA probe or a scrambled control detection probe (green) and visualized using FISH. miR-124 is ubiquitously expressed in blastula and gastrula stages and is enriched in the ciliary band of the larvae. 3 biological replicates. Scale bar=50 µm. (B) Double FISH is performed against DIG labeled miR-124 LNA (green) and fluorescence labeled *Onecut* (*Hnf6)* (magenta) and counterstained with DAPI (blue). 2 biological replicates. Maximum intensity projection of Z-stack confocal images is presented.

### Inhibition of miR-124 leads to developmental defects

To test the loss-of-function of miR-124, we microinjected miR-124 Locked-Nucleic-Acid (LNA) inhibitor into zygotes. Embryos injected with miR-124 inhibitor have a statistically significant reduction of miR-124 levels, compared to embryos injected with the Texas Red dextran, indicating the effectiveness of the inhibitor (Fig. 3A). We observed a dose-dependent severity of miR-124 inhibitor induced phenotypes, ranging from a developmental delay, gut morphological defects, lack of visible coelomic pouches (which contain the multipotent stem cells), clusters of cells in the blastocoel of gastrula stage embryos, and combinations of these defects (Figs. 3B, 3C). Of note is that coelomic pouches are present at the larvae stage (72hpf), suggesting a transient delay in their formation (Fig. S3).

**Figure 3.**
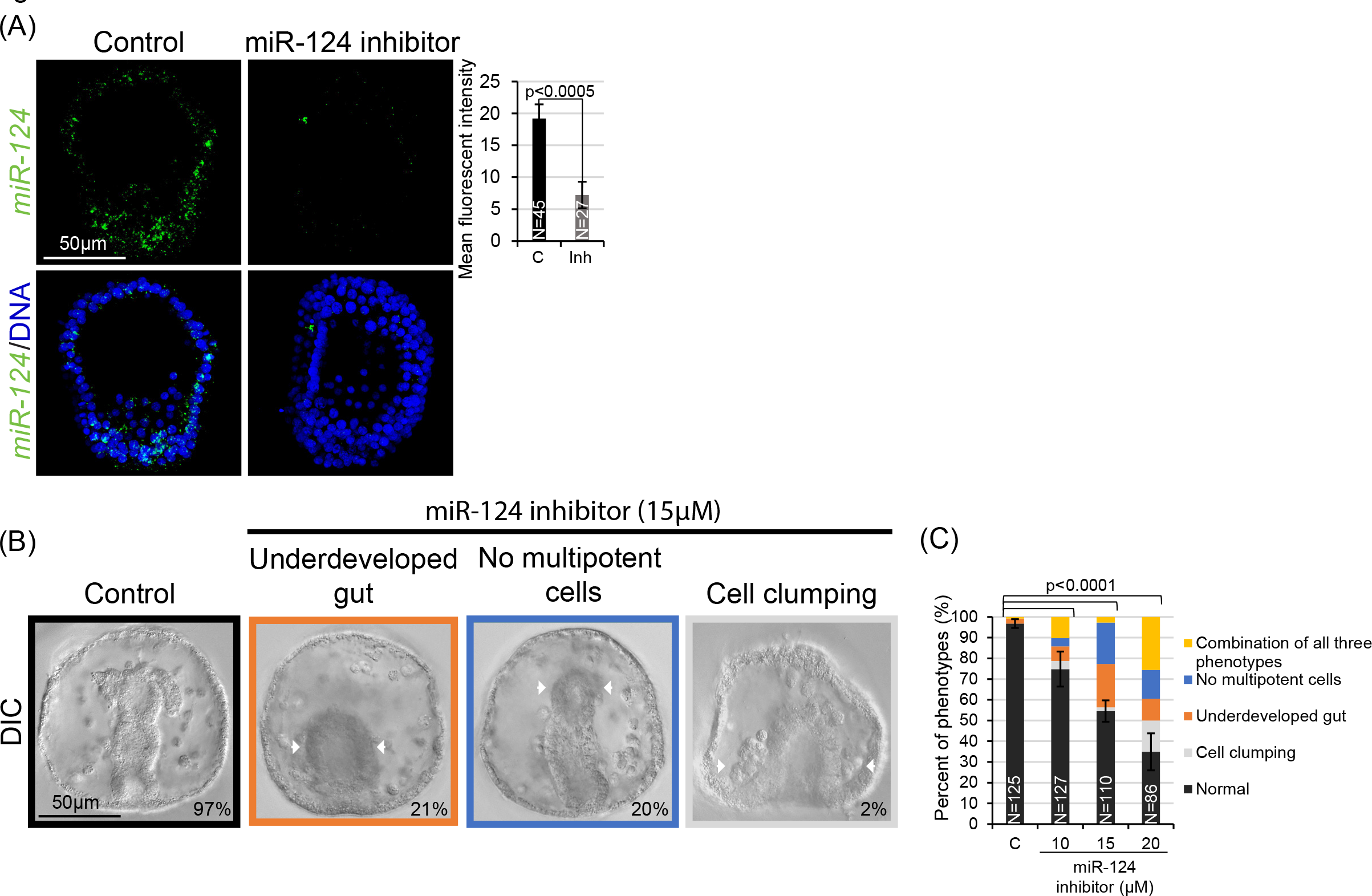
miR-124 inhibition results in dose-dependent developmental defects. (A) miR-124 LNA power inhibitor was injected into zygotes and cultured to 24 hpf, followed by miR-124 FISH (green). Embryos were counterstained with DAPI to label DNA (blue). miR-124 level was greatly reduced in the presence of the miR-124 inhibitor. Scale bar=50 µm. Maximum intensity projection of Z-stack confocal images is presented. Average fluorescent intensity was calculated. Student t-test was used. 3 biological replicates. (B) Developmental defects in miR-124 inhibitor injected embryos, including underdeveloped gut, undetectable coelomic pouches, and cell clumping. Defects are indicated by the white arrows. (C) The severity of the defects was observed in a dose- dependent manner. 4 biological replicates. C=Control, Inh= miR-124 inhibitor injected embryos, N= total number of embryos. For all the graphs SEM is graphed. Cochran- Mantel-Haenszel statistical test was used.

### Inhibition of miR-124 results in decreased gut contractions

One of the morphological changes we observed in miR-124 inhibitor injected gastrulae is having a significantly wider gut compared to the control (Fig. 4A). We further examined the gut tissues at the molecular level and found that the miR-124 inhibitor injected gastrulae failed to express Endo1, which is expressed specifically in the midgut and hindgut (Fig. 4A)^73^. These gut defects were rescued with a co-injection of miR-124 inhibitor and a miR-124 mimic, indicating that these defects are specifically due to the miR-124 inhibition (Fig. 4A). Interestingly, by the larval stage, these miR-124 inhibitor injected larvae express similar levels of Endo1 as the control larvae, indicating that that lack of Endo1 expression in the gastrulae is a transient defect (Fig. 4B). In addition, we found that miR-124 inhibition resulted in decreased gut contractions compared to the control in the larvae (Fig. 4C, Video 1). To elucidate potential mechanisms of the gut defects, we examined the expression of *FoxA* and *Krl*, which are key transcription factors important for endodermal specification^74, 75^. Results indicate that the expression of *FoxA* and *Krl* did not change significantly (Fig. S4). Next, we examined if changes in gut contractions were due to defects in the mesodermally-derived muscle. Results indicate that filamentous actin (F-actin) of the gut circumpharyngeal muscles were similarly structured in miR-124 inhibitor injected larvae and control. However, we observed that miR-124 inhibitor-injected larvae had a significantly wider cardiac sphincter, which separates the fore- and midgut of the embryo, in comparison to the control larvae (Fig. 4D)^76–78^. Thus, defects in gut contractions of miR-124 inhibitor injected larvae may be partially due to the cardiac sphincter defects and/or neuronal network of the gut. The mechanism of how miR-124 inhibition induced these gut defects still needs to be elucidated.

**Figure 4.**
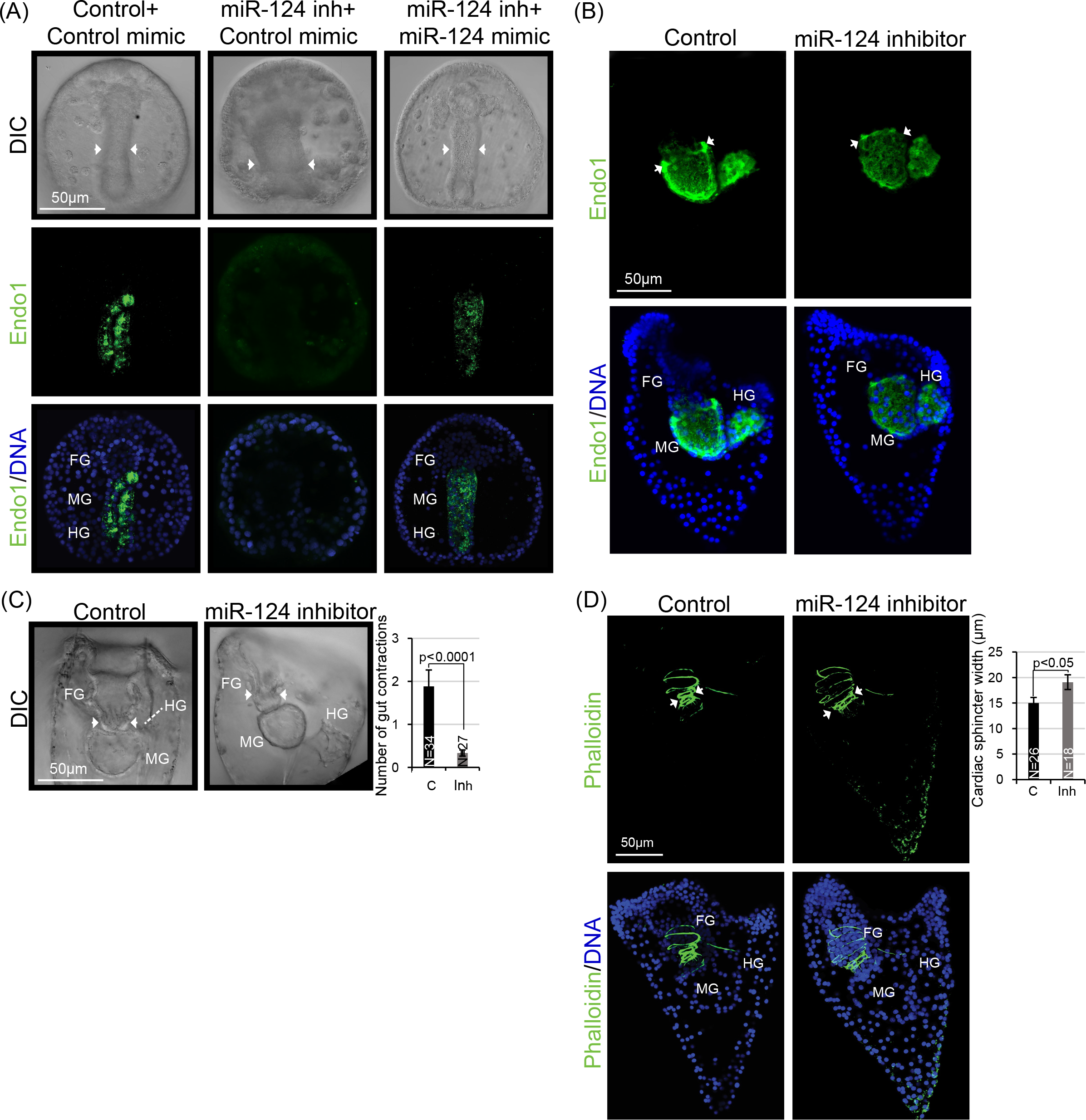
miR-124 inhibition leads to gut and sphincter defects. (A) miR-124 inhibitor injected gastrulae were collected at 48 hpf and immunolabeled with Endo1 (green) which detects antigen of the mid- and hindgut. miR-124 inhibitor injected gastrulae exhibited a significantly wider gut and failed to express endodermal antigens (Endo1). These defects were rescued with a co-injection of the miR-124 inhibitor and a miR-124 mimic. Embryos were counterstained with DAPI (blue). 3 biological replicates. FG=Foregut, MG=Midgut, HG=Hindgut. Control N=12, miR-124 inhibitor injected embryos N=10, and miR-124 inhibitor+miR-124 mimic N=7. White arrows point to the width of the midgut. (B) Larvae were immunolabeled with Endo1 antibody and counterstained with DAPI. 3 biological replicates N=10. (C) miR-124 inhibitor injected larvae exhibited a significant decrease in gut contractions over a four-minute period. images are still frames of embryos during gut contractions. 3 biological replicates. (D) Phalloidin stain (green) indicated that miR-124 inhibitor injected larvae had wider cardiac sphincter compared to controls. Embryos were counterstained with DAPI (blue). FG=Foregut, MG=Midgut, and HG=Hindgut. White arrows delineate the width of the cardiac sphincter. 3 biological replicates. SEM is graphed. Student t-test was used. N= total number of embryos. C=Control. Inh=miR-124 inhibitor injected embryos. Maximum intensity projection of Z-stack confocal images is presented.

### miR-124 regulates larval swimming

Sea urchin larval swimming is driven by force generated by beating of the cilia on the ciliary band and is regulated by several neurotransmitters, including serotonin, dopamine, and γ-aminobutyric acid^22, 31, 79–81^. We observed a significant decrease in the swimming velocity (distance/time) in the miR-124 inhibitor injected larvae compared to the control (Fig. 5A, Video 2). Larvae co-injection with the miR-124 inhibitor and miR- 124 mimic have restored swimming compared to larvae injected with miR-124 inhibitor: miR-124 inhibitor and miR-124 mimic injected larvae have their swimming velocity 17% less than the control, compared to miR-124 inhibitor injected embryos that have their swimming velocity 73% less of the control embryos (Fig. 5A, Video 2). These results indicate that this swimming defect is specifically induced by miR-124 inhibition.

**Figure 5.**
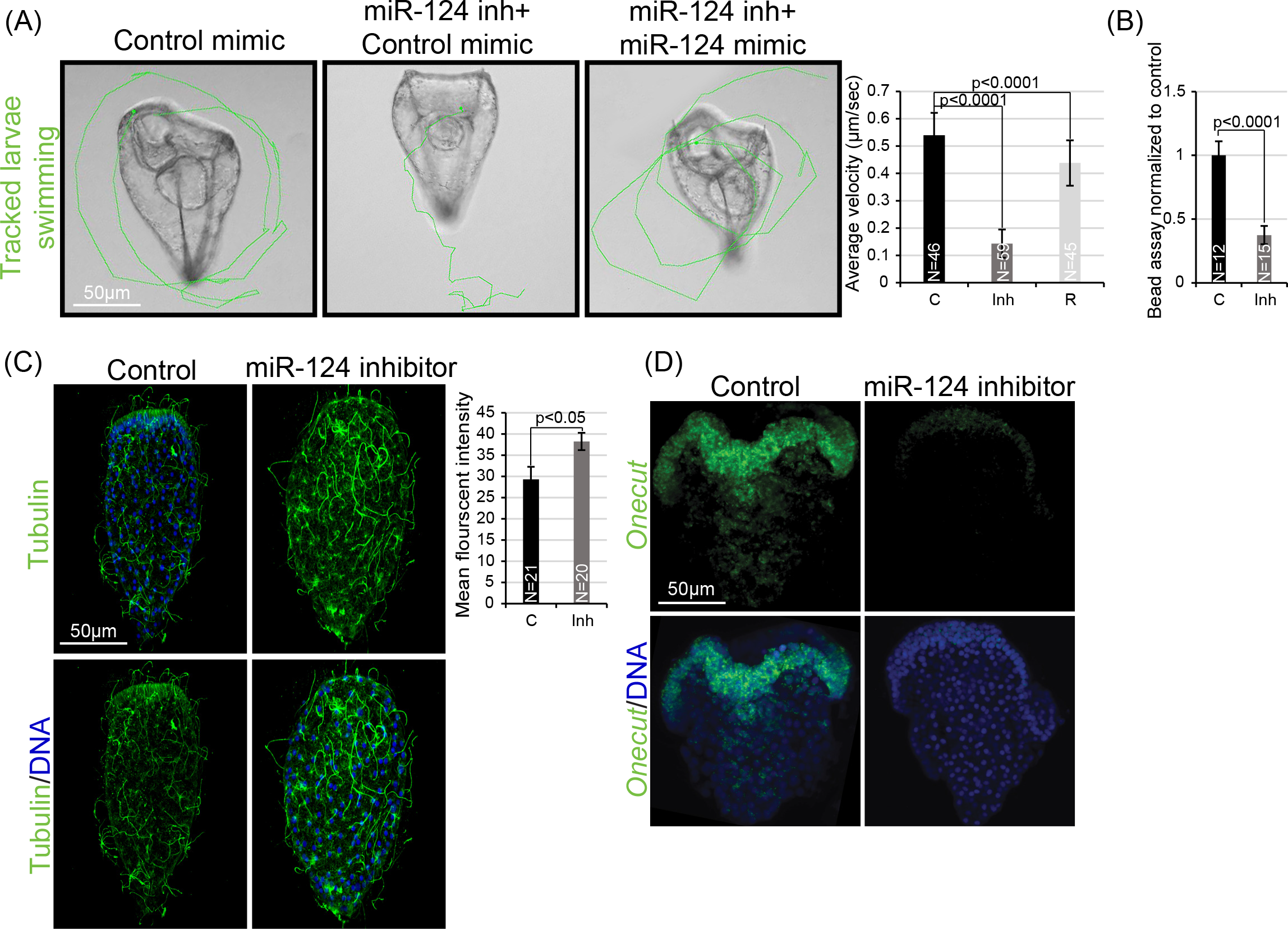
miR-124 inhibition results in decreased larval swimming velocity. (A) Larvae were imaged live for 60 seconds and tracked manually with ImageJ plugin to obtain the velocity. miR-124 inhibitor injected embryos exhibited a significant decrease in swimming velocity compared to the control. These defects were rescued with a co-injection of miR-124 inhibitor and miR-124 mimic. 3 biological replicates. Tracks of larvae swimming path is depicted in green. (B) Embryos were imaged lived for cilia beating for 120 seconds with polybeads. The average number of beads that flowed around the control embryo was significantly higher compared to the miR-124 inhibitor injected larvae. 3 biological replicates. (C) The cilia of control and miR-124 inhibitor injected embryos were immunolabeled with tubulin antibody (green). miR-124 inhibitor injected larvae exhibited an increase in fluorescent intensity in tubulin compared to the control. Control embryos were injected with a non-fixable FITC dextran and the miR-124 inhibitor injected embryos were coinjected with fixable Texas Red dextran. These embryos were incubated in the same well during immunolabeling and imaging. Larvae were counterstained with DAPI to label DNA (blue). 3 biological replicates. SEM is graphed. Student t-test was used. (D) miR-124 inhibitor injected larvae exhibited a decrease in *Onecut (Hnf6)* expression compared to the control. 4 biological replicates. C=51 and miR-124 inhibitor injected embryos=52.

We further assessed the effectiveness of ciliary beating by counting the number of polybeads propelled by the larval ciliary beating in the anterior region of the control or miR-124 inhibitor injected larvae (Fig. 5B, Video3). Results indicate that control embryos propelled a greater number of beads compared to the miR-124 inhibitor injected embryo, indicating that miR-124 inhibitor injected larvae are less effective at coordinating ciliary beating (Fig. 5B, Video 3). Using tubulin immunolabeling, we did not observe differences in the morphology of the cilia between the miR-124 inhibitor injected and the control larvae. However, a significant increase in tubulin levels was consistently observed in the miR-124 inhibitor injected larvae compared to the control (Fig. 5C).

Since swimming is controlled by ciliary band beating, the proper formation of the ciliary band is important. The expression of *Onecut* has been shown to play a role in ciliary band formation, although the mechanism of its function is still unknown^82^. We observed that the expression of *Onecut* was dramatically decreased in miR-124 inhibitor injected larvae compared to the control (Fig. 5D). This decrease in *Onecut* expression can potentially negatively affect the formation of the layered epithelial cells of the ciliary band which is important for the formation of neuronal connections^81, 83, 84^.

### Inhibition of miR-124 leads to decreased mature neurons

Serotonin has been found to mediate sea urchin gut contractions and larval swimming behavior^31, 32, 79, 80, 85, 86^. Since we observed gut contraction and swimming defects, we examined serotonin and found that while the number of serotonergic neurons stayed the same in miR-124 inhibitor injected larvae compared to the control, the overall level of serotonin in the miR124 inhibitor injected larvae was significantly decreased along with less dendritic spines compared to the control (Fig. 6A). We also examined the mature neuronal network with the sea urchin neuronal antibody (1E11 and SynB) which recognizes neuron-specific SynB in the larval mouth, apical organ, ciliary band, and neurites projecting from the ciliary band^26, 32, 87^. miR-124 inhibition resulted in a significant decrease in SynB expressing neurons along the ciliary band and the mouth (Fig. 6B). This decrease in mature neurons was rescued with a co-injection of miR-124 inhibitor with miR-124 mimic, indicating that the observed neuronal defects are specifically induced by the miR-124 inhibitor (Fig. 6B).

**Figure 6.**
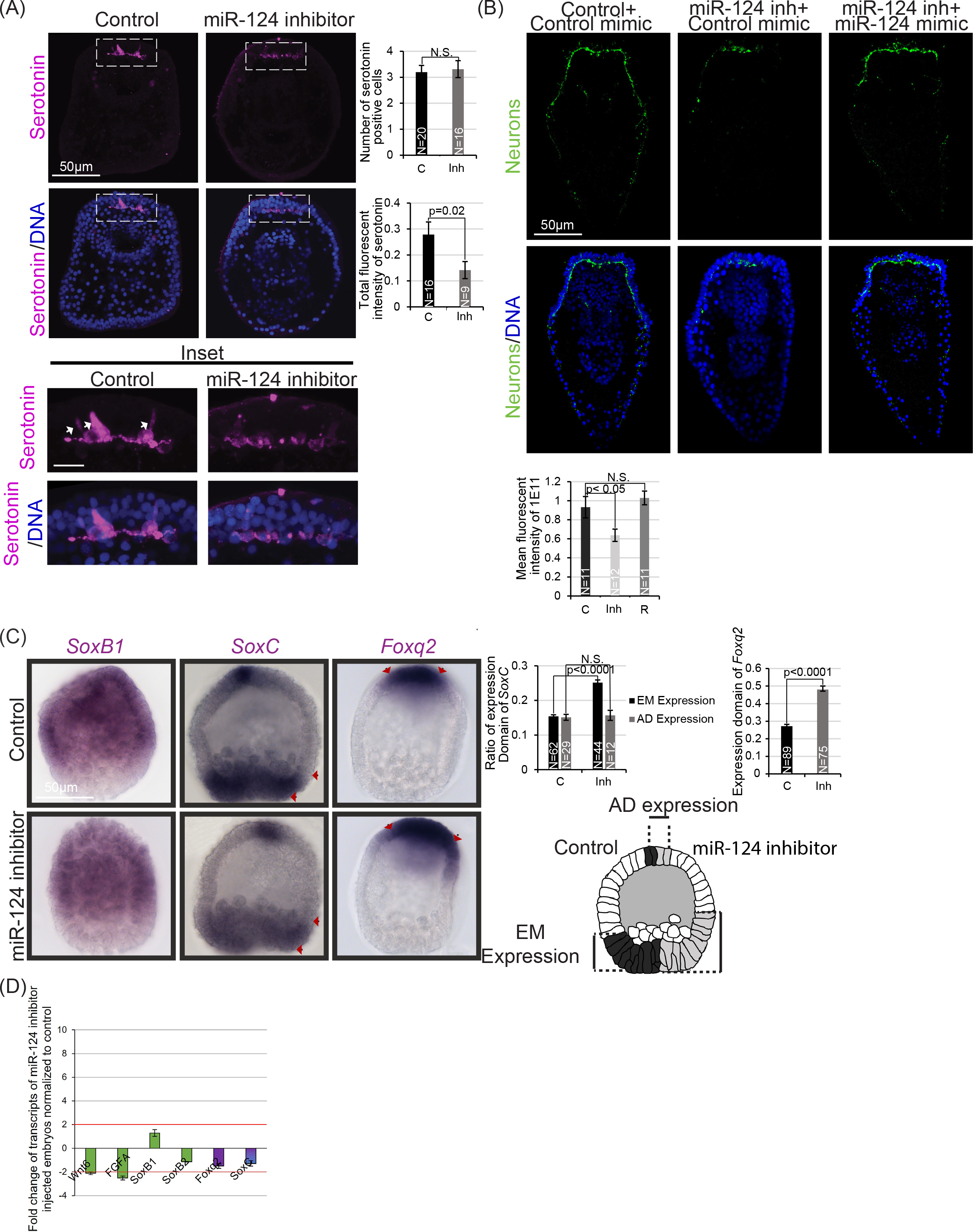
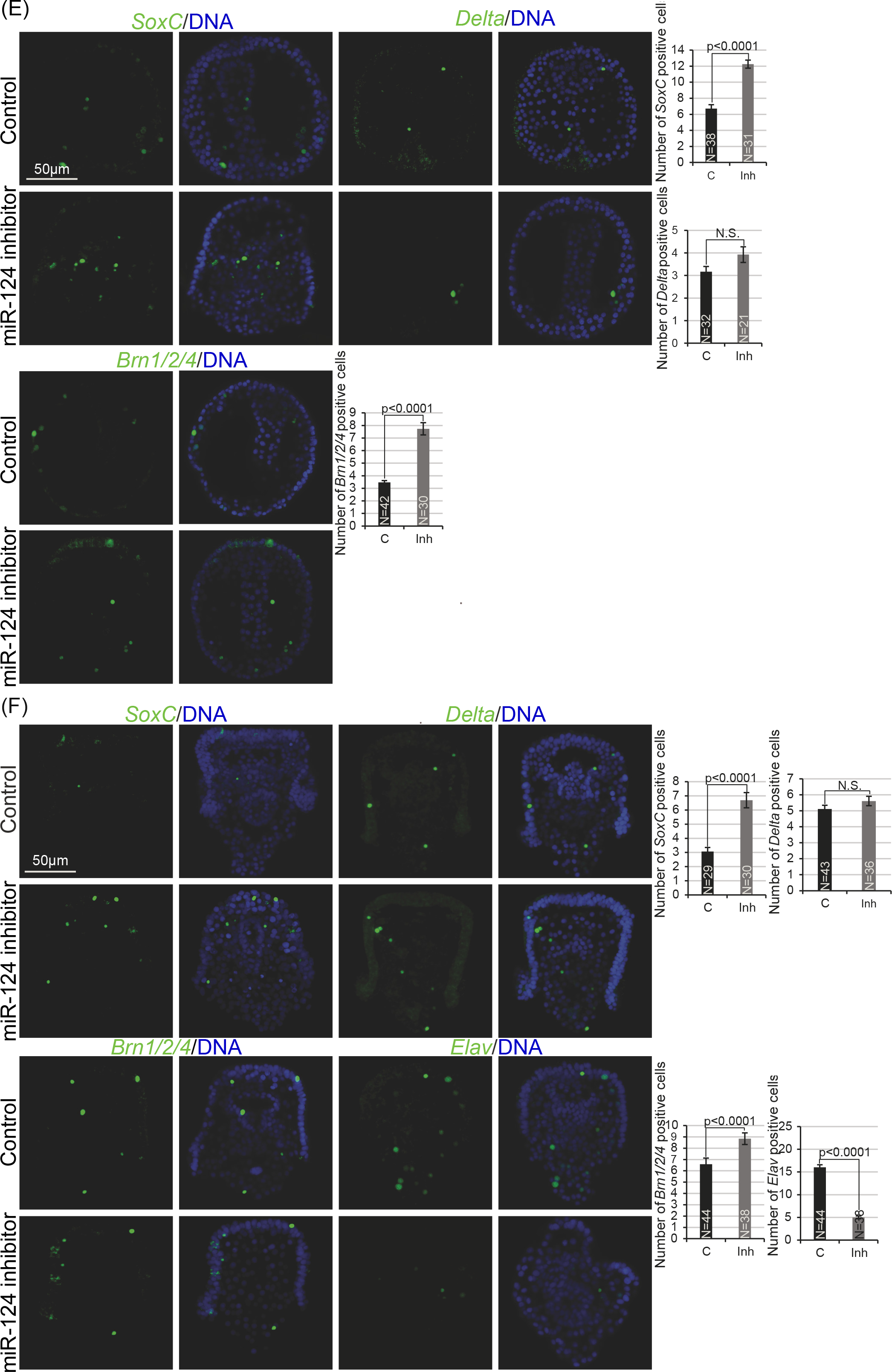
miR-124 regulates neurogenesis. (A) Oral view of serotonin-containing neurons (magenta) with close-up view (shown by the inset delineated by the white dashed lines). Larvae were immunolabeled with the serotonin antibody (magenta) and counterstained with DAPI (blue). miR-124 inhibitor injected larvae had significant reduction in levels of serotonin compared to the control. The number of serotonin positive cells did not differ. 3 biological replicates. (B) miR-124 inhibitor injected larvae were immunolabeled with sea urchin neuronal antibody 1E11 that detects SynB expressing neurons (green) and counterstained with DAPI (blue). miR-124 inhibitor injected larvae exhibited a decrease in SynB expressing neurons. Co-injection of miR- 124 inhibitor with miR-124 mimic resulted in a recuse of the decrease in SynB expressing neurons. 3 biological replicates. Scale bar=50 µm. Maximum intensity projection of Z-stack confocal images is presented. (C) *SoxB1, Foxq2*, and *SoxC,* expression domains at the blastula stage was significantly increase in miR-124 inhibitor injected embryos. 3 biological replicates. (D) 100 of control and miR-124 inhibitor injected embryos were collected at the blastula stage and neuronal transcripts were assessed. Wnt6 and FGFA were 2-fold decreased in miR-124 inhibitor injected embryos compared to the control. (E) *SoxC* and *Brn1/2/4* expression was increased in the miR- 124 inhibitor injected gastrulae compared to the control, while *Delta* expression did not change (green). Embryos were counterstained with DAPI (blue). 3 biological replicates. (F) The numbers of *SoxC, Delta,* and *Brn1/2/4* expressing cells (green) were increased in miR-124 inhibitor injected larvae compared to the control, while the number of *Elav* positive cells was decreased in miR-124 inhibitor injected larvae compared to the control. Larvae were counterstained with DAPI to label DNA (blue). 3 biological replicates. EM=Endomesoderm. AD=Apical domain. C=Control. KD=miR-124 inhibitor injected embryos. N= total number of embryos. SEM is graphed. Student’s t-test was used. Maximum intensity projection of Z-stack confocal images is presented.

### miR-124 modulates neuronal GRN to regulate specification, differentiation, and maturation of neurons

To systematically examine the function of miR-124 on the specification, differentiation, and maturation of neurons, we tested the spatial and temporal expression of neuronal GRN components in control and miR-124 inhibitor injected embryos. *Wnt6* and *FGFA* transcripts were 2-fold decreased in miR-124 inhibitor injected embryos compared to the control (Fig. 6D). SoxB1 is a TF at the top of the neuronal GRN hierarchy, in regulating all three domains of the nervous system^23, 88, 89^. In the miR-124 inhibitor injected blastulae, *SoxB1* expression was not different from the control (Fig. 6C, 6D). Downstream of SoxB1 is Foxq2, which is important for establishing the neuronal apical domain in the early embryo^22, 88^. The lack of *Foxq2* expression later in gastrula stage (48 hpf) is important for proper development of serotonin expressing neurons in the ciliary band^29, 30^. A significant expansion of *Foxq2* expression is observed in the miR-124 inhibitor injected embryos compared to the control (Fig. 6C); however, the level of *Foxq2* transcripts is not significantly changed (Fig. 6D). Downstream of Foxq2 is SoxC, which is important for development of neurons in the apical domain as well as the endomesoderm^81, 90, 91^. SoxC and Foxq2 expression regulates neuronal progenitor specification in the sea urchin apical domain during the blastula stage^21, 22, 34^, consistent with role of *SoxC* in progenitor neurons in other systems^28, 35, 81, 92^. The miR-124 inhibitor injected larvae had a significant increase in *SoxC* expression in the endomesoderm region compared to the control at the blastula stage but did not change in the apical domain, suggesting that SoxC may have additional function in the embryo^15, 89, 93^ (Fig. 6C). Overall, the level of *SoxC* transcripts was not significantly altered by miR-124 perturbation at the blastula stage (Fig. 6D).

Later, the *SoxC*-positive neuronal progenitor cells also express *Brn1/2/4*^22^. miR-124 inhibitor injected gastrulae have increased number of *SoxC* and *Brn1/2/4* positive cells (Fig. 6E). During gastrula to larval stages, a small number of cells express *Delta* to mediate differentiation of neurons during the larval stage^24, 90, 94, 95^. Once the neuronal *SoxC*-positive progenitors undergo their last mitotic division mediated by Delta/Notch signaling, the differentiated neuron becomes *Brn1/2/4* and *Delta* positive. We did not observe any change in the number of *Delta* expressing cells at the gastrula stage (Fig. 6E). However, miR-124 inhibition resulted in an increase of *SoxC* and *Brn1/2/4* positive cells in the gastrula stage that persisted to the larval stage, indicating that miR-124 has a broad impact on several TFs regulating the neuronal differentiation process (Fig. 6C- F). We further tested the number of *Elav* expressing cells to examine neuronal maturation. Elav is an RNA binding protein that is known to stabilize transcripts important for axonal targeting and synaptic growth^19, 22^. Elav protein is expressed in cells throughout the cytoplasm and in differentiated neurons within the ciliary band and apical domain of the sea urchin larvae^22^. We observed that the number of *Elav* expressing cells is significantly lower in the miR-124 inhibitor injected larvae compared to the control (Fig. 6F). Overall, these results indicate that miR-124 inhibition led to an increased number of cells expressing neuronal specification and differentiation factors and a concomitant decrease of mature and functional neurons.

### miR-124 directly suppresses components of neuronal GRN

To determine the molecular mechanism of miR-124’s regulation of neurogenesis and larval behavior, we performed a bioinformatic analysis and identified *NeuroD1* and *Notch* as potential miR-124 targets. We cloned the *NeuroD1* or *Notch*, downstream of *Renilla Luciferase* (*RLuc*) reporter construct. Site-directed mutagenesis was used to disrupt miR-124 seed sequences within the *NeuroD1* and *Notch*. *Firefly* (*FF*) luciferase flanked by *β-globin* UTRs was used as an injection control. The dual luciferase assay results indicated that miR-124 directly suppresses *NeuroD1* and the first seed site within *Notch* (Fig. 7A).

**Figure 7.**
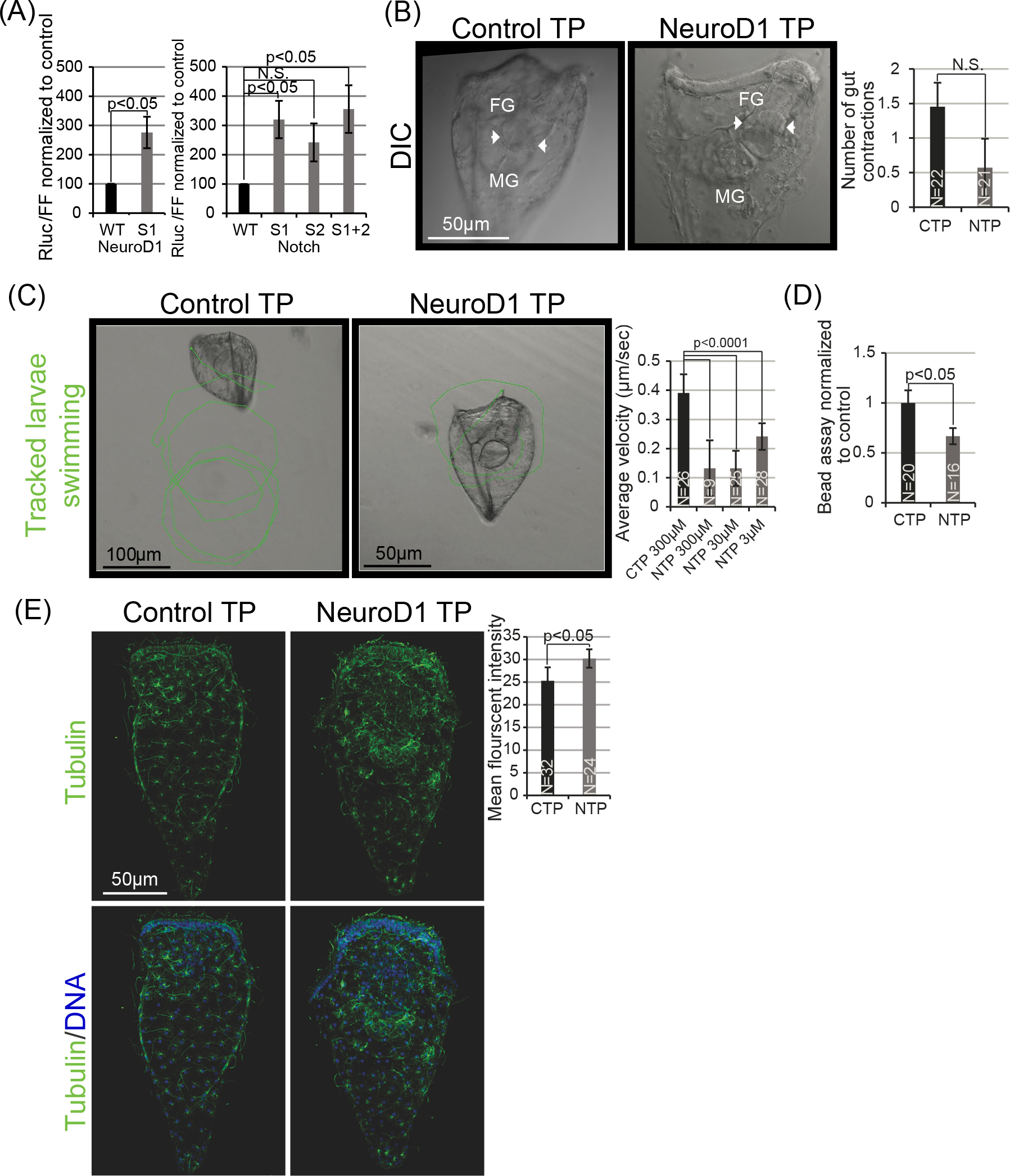
miR-124 directly suppresses *NeuroD1* to regulate swimming. (A) *Notch* and *NeuroD1* dual luciferase assays indicated that miR-124 directly suppresses *Notch* at seed1 and *NeuroD1* seed. Each biological replicate contained 50 embryos. 3 biological replicates. (B) Removing miR-124 suppression of *NeuroD1* using *NeuroD1* TP resulted in decreased but not significant gut contractions in a span of 4 minutes. Representative images of gut contractions are depicted. (D) Larvae were imaged live and imaged for 60 seconds and manually tracked with ImageJ plugin to obtain the velocity. *NeuroD1* TP larvae exhibited a significant decrease in swimming velocity compared to the control. 3 biological replicates. Tracks of larvae swimming are depicted in green. (E) Embryos were imaged lived for cilia beating for 60 seconds. 3 biological replicates. (F) *NeuroD1* TP larvae exhibited an increase in tubulin (green) compared to the control. Larvae were counterstained with DAPI to label DNA (blue). 3 biological replicates. For all the graphs SEM is graphed. Student t-test was used.

### Removing miR-124’s direct suppression *NeuroD1* results in swimming defects

Since the regulatory role of the Delta/Notch signaling pathway and miR-124’s regulation of *Notch* on neuronal development have been examined previously^24, 58, 90^, we focus here on examining miR-124’s regulation of *NeuroD1*. To determine the impact of removing miR-124 suppression of *NeuroD1*, we injected a morpholino-based target protector (TP)^96, 97^ that is complementary to the miR-124 binding site and flanking sequences within *NeuroD1*. This *NeuroD1* TP will prevent the endogenous miR-124 from binding to the *NeuroD1* 3’UTR to mediate post-transcriptional repression.

Removing miR-124 suppression of *NeuroD1* resulted in a trend of decreased gut contractions (Fig. 7B). When we injected zygotes with exogenous *NeuroD1* transcripts to mimic the effect of blocking miR-124’s suppression of *NeuroD1*, we observed that gut contractions were significantly decreased, indicating that *NeuroD1* overexpression (OE) is sufficient to result in aberrant gut contractions (Fig. S5A, Video 7).

For swimming behavior, results indicate that *NeuroD1* TP injected zygotes displayed similar swimming defects as miR-124 inhibitor injected larvae and *NeuroD1* OE larvae (Fig. 7C, Video 4; 5A, Video 5; Fig. S5B, Video 8). *NeuroD1* TP injected larvae also exhibited decreased efficacy in cilia beating, as assayed by the larvae’s ability to propel beads (Fig. 7D, Video 6). Removal of miR-124’s suppression of *NeuroD1* resulted in a slight increase in tubulin, similar to what was observed in miR- 124 inhibitor injected larvae (Fig. 7E).

### Removing miR-124 suppression of *NeuroD1* results in decreased functional neurons and increased *Elav* expressing cells

Similar to miR-124 inhibitor injected larvae, *NeuroD1* TP injected embryos had a significant decrease in the overall level of serotonin, while the number of serotonin expressing cells stayed the same compared to the control (Fig. 8A). Decrease in serotonin levels was also observed in *NeuroD1* OE larvae (Fig. S5C). In addition, removing miR-124 suppression of *NeuroD1* resulted in a significant decrease in SynB- expressing neurons along the ciliary band and the mouth (Fig. 8B). This change in SynB-positive neurons was also observed in the *NeuroD1* OE larvae (Fig. S5D).

**Figure 8.**
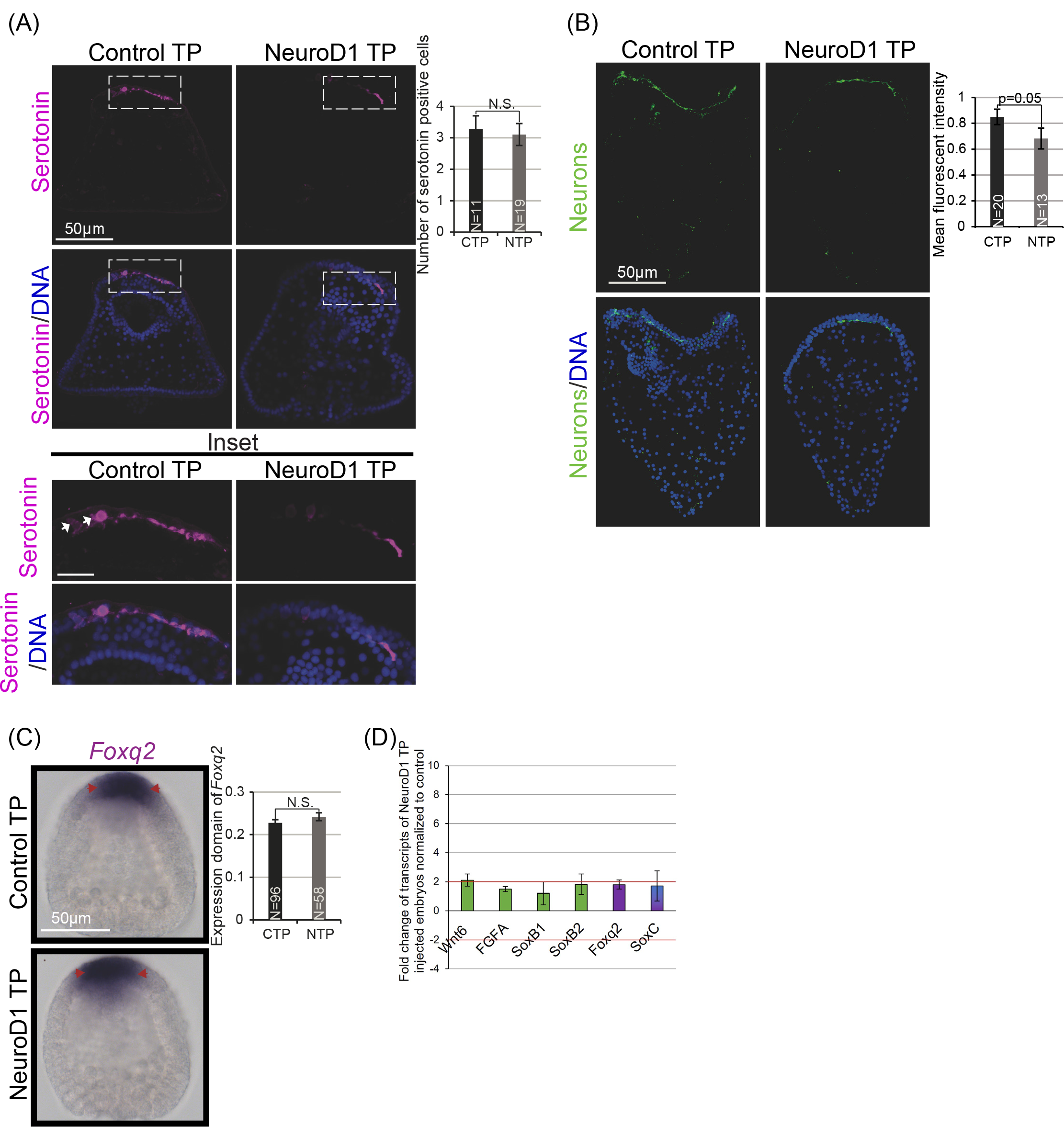
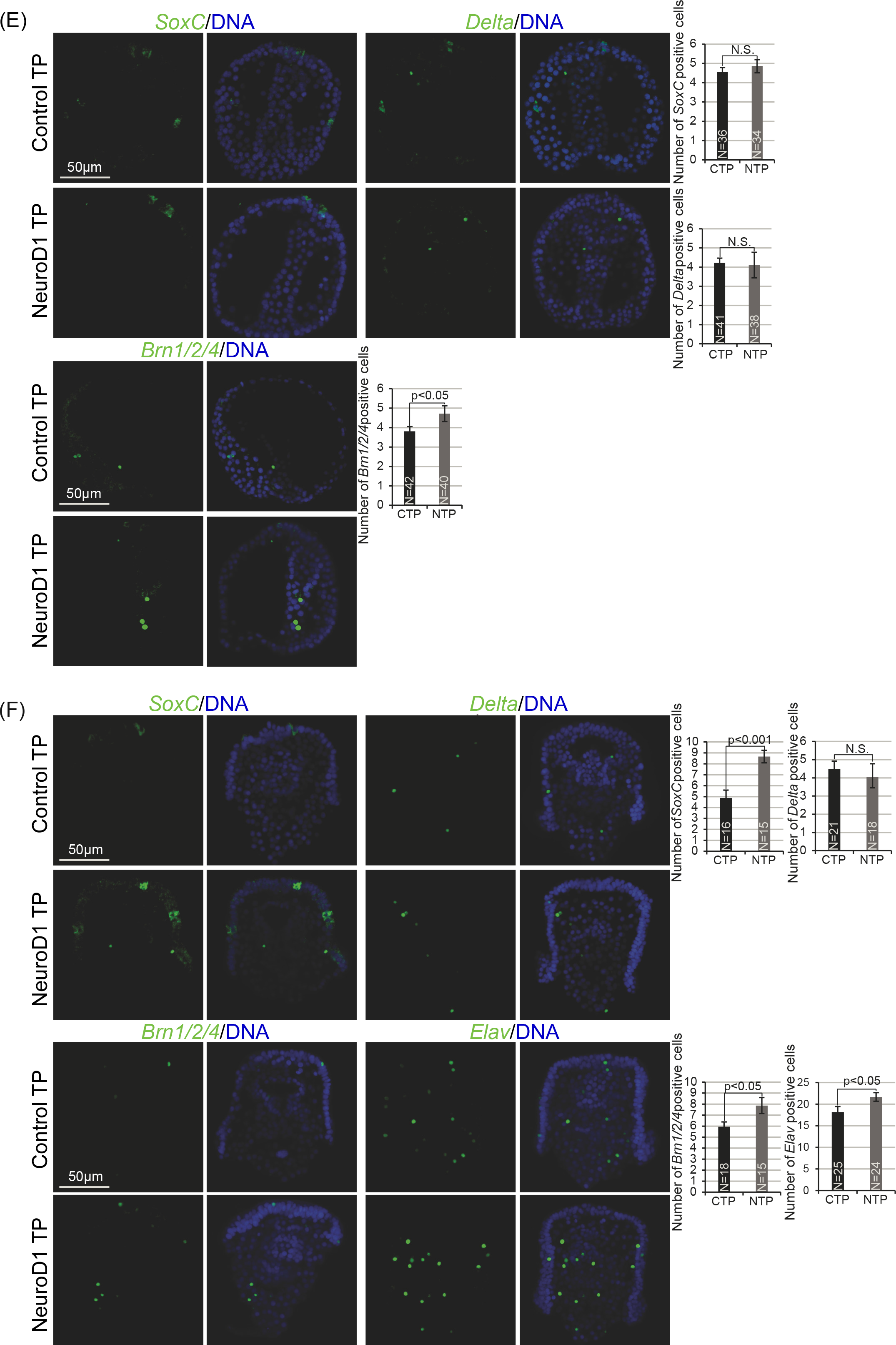
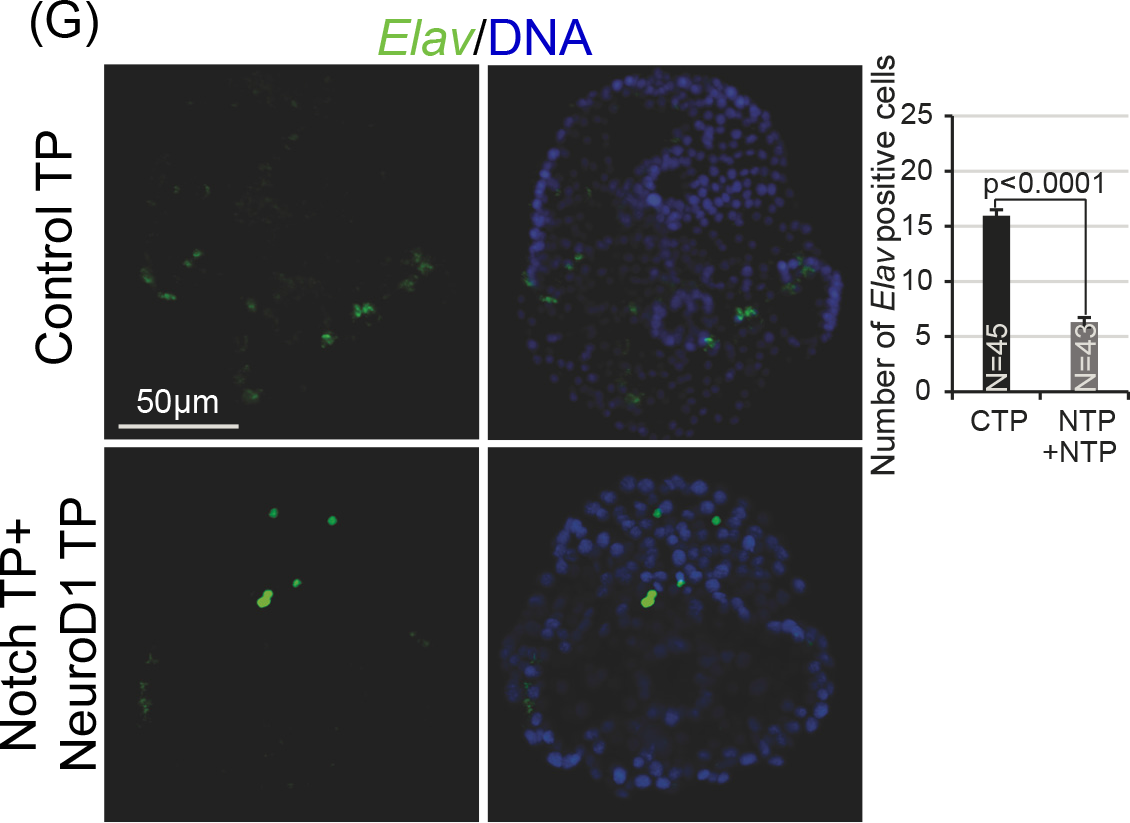
miR-124’s direct regulation of NeuroD1 is important for neurogenesis. (A) Oral view of serotonin-containing neurons (magenta) with close-up view (shown by the inset delineated by the white dashed lines). Larvae were immunolabeled with the serotonin antibody (magenta) and counterstained with DAPI (blue). 3 biological replicates. (B) *NeuroD1* TP larvae were immunolabeled with sea urchin neural antibody 1E11 that detects SynB expressing neurons (green) and counterstained with DAPI (blue). *NeuroD1* TP larvae exhibited significant decrease in mature neurons. 3 biological replicates. Maximum intensity projection of Z-stack confocal images is presented. (C) *SoxB1* and *Foxq2* expression domains were similar in control and *NeuroD1* TP injected blastulae. 3 biological replicates. (D) 100 control or *NeuroD1* TP injected embryos were collected for qPCR and assayed for changes in neuronal transcripts. (E) The number of *Brn1/2/4* expressing cells was increased in the *NeuroD1* TP injected gastrulae compared to the control. 3 biological replicates. (F) The numbers of *SoxC, Brn1/2/4,* and *Elav* expressing cells (green) were increased in the *NeuroD1* TP injected larvae compared to the control. Larvae were counterstained with DAPI to label DNA (blue). 3 biological replicates. CTP=Control Target Protector. NTP=NeuroD1 Target Protector. EM=Endomesoderm. AD=Apical domain. N= total number of embryos. SEM is graphed. Student’s t-test was used. Maximum intensity projection of Z-stack confocal images is presented. (G) *NeuroD1* TP was co-injected with *Notch* TP at 30µM each with corresponding control. A significant decrease in *Elav-*expressing cells was observed in larvae coinjected with *NeuroD1* TP and *Notch* TP compared to the control.

To further reveal the impact of miR-124’s suppression of *NeuroD1*, we systematically examined the spatial expression of factors of the neuronal GRN. The spatial expression of *Foxq2* was similar between *NeuroD1* TP or control TP-injected blastulae (Fig. 8C). No significant change in transcript levels of all the major neuronal factors in control or *NeuroD1* TP injected blastulae was observed (Fig. 8D).

In the gastrula stage, *NeuroD1* TP injected embryos exhibited no change in the number of *SoxC* and *Delta* expressing cells and had consistently one additional Brn1/2/4 expressing cell compared to the control (Fig. 8E). However, in the larval stage, *NeuroD1* TP injected embryos have a significant increase in *SoxC, Brn1/2/4*, and *Elav* positive cells compared to the control (Fig. 8F).

### miR-124 regulates *NeuroD1* and *Notch* in the neuronal GRN

The increased number of *Elav* expressing cells in *NeuroD1* TP injected larvae contrasts with the decreased number of *Elav* expressing cells in miR-124 inhibitor injected embryos (Figs. 6F and 8F). To resolve the difference of the number of *Elav* expressing cells between *NeuroD1* TP and miR-124 inhibitor injected embryos, we co-injected *NeuroD1* TP with *Notch* TP to test the effects of removing miR-124’s suppression of both transcripts and assay for changes in the number of *Elav* expressing cells. Results indicate that co-injection of *NeuroD1* TP and *Notch* TP recapitulated the decrease in *Elav* expressing cells observed in miR-124 inhibitor injected embryos (Figs. 6F and 8G). This result suggests that miR-124’s suppression of NeuroD1 resulted in increased numbers of mature *Elav* expressing cells. However, miR-124’s suppression of *Notch* may promote neuronal progenitor cells to undergo excessive Delta/Notch mediated apoptosis that leave fewer differentiated neurons in miR-124 inhibitor injected or in coinjected *NeuroD1* TP and *Notch* TP larvae compared to the control.

## Discussion

We integrated NeuroD1 into the neuronal gene regulatory network and identified that it regulates *SoxC*, *Brn1/2/4*, *Delta* and *Elav*. With a more complete neuronal GRN, we systematically examined the post-transcriptional regulation mediated by miR-124.

We discovered that miR-124 regulates gut contractions, swimming behavior, and neuronal development. Some of these miR-124 perturbation induced phenotypes may be attributed to miR-124’s direct suppression of *Notch* and *NeuroD1* that resulted in decreased serotonin and functional neurons. The molecular mechanism of miR-124’s regulation of neurogenesis is in part through its suppression of *Notch*, which mediates the final mitotic division of progenitor neurons to differentiated neurons^15, 22, 26^. Further, miR-124 also suppresses *NeuroD1*, which we find to be important in mediating the transition between differentiation and maturation of neuronal development. Overall, this study contributes to our understanding of miR-124’s prolific regulatory role in neuronal specification, differentiation, and maturation.

Previously, it has been observed that *NeuroD1* is expressed in the larval gut and ciliary band^93^, consistent with our finding that NeuroD1 protein is present in the presumptive ganglia and neurofilaments as well as in the gut (Fig. 1C). In vertebrate systems, in addition to its function in neurogenesis, NeuroD1’s loss-of-function studies revealed that NeuroD1 has additional function in the pancreas, where its loss resulted in severe diabetes^46^. Interestingly, cells with a pancreatic-like signature are localized within the sea urchin embryonic gut and that pancreatic endocrine cells and neurons express similar transcription factors during development^93, 98^. It was proposed that these pancreatic endocrine cells in the larval gut may have co-opted some neuronal regulatory factors from an ancestral neuron^93, 98^.

In the *NeuroD1* knockdown gastrulae, expression of *Delta* is increased 2.5-fold, suggesting that NeuroD1 may potentially repress *Delta* in the gastrula to commit progenitor cells to adopt the neuronal fate, and/or NeuroD1 may repress *Delta* to regulate additional non-neuronal processes (Fig. 1D). In the larval stage, NeuroD1 may perform its activation of neuronal factors, including *SoxC*, *Brn1/2/4*, *Delta*, and *Elav* (Fig. 1D). Of these TFs, *SoxC*, *Brn1/2/4*, and *NeuroD1* transcripts have also been found to be expressed in the apical domain and ciliary band of the sea urchin larvae, as well as in the gut, indicating that NeuroD1 plays a conserved role in promoting neuronal differentiation^50, 51, 53, 71^ and potentially additional processes in the endomesoderm. How NeuroD1 temporally regulates the expression of *Delta* during gastrula and larval stages of development needs to be investigated further.

To understand the function of miR-124, we injected miR-124 inhibitor into zygotes. One of the defects we observed was decreased gut contractions in miR-124 inhibitor injected larvae compared to the control (Fig. 4C). A potential explanation may be due to decreased level of serotonin (Fig. 6A), as it has been shown that serotonin binds to the receptors near the pyloric sphincter to allow for sphincter opening^42^. Since we observe a trend in decreased gut contractions in the *NeuroD1* TP injected larvae, whereas miR-124 inhibition and *NeuroD1* overexpression resulted in significant decrease in gut contractions compared to the control, this suggests that miR-124 is likely to regulate NeuroD1 and an additional unknown factor to impact gut contractions (Figs. 4C, 7B, and S5A). The exact molecular mechanism of how miR-124 regulates gut development and gut contractions is still unknown.

Even though miR-124 has a conserved role in neurogenesis^52, 56–59, 99, 100^, little research has been conducted on miR-124’s regulation of serotonin^55, 101^. We observed no change in the number of serotonin positive cells between miR-124 inhibitor injected embryos and control embryos, suggesting that the specification of these cells is not affected by miR-124 perturbation. However, the overall level of serotonin was significantly decreased in the miR-124 inhibitor injected embryos compared to the control (Fig. 6A). Serotonin is a neurotransmitter important for early swimming, feeding behavior, and gut contraction in the larvae embryo^31, 32^. Decreased serotonin level in miR-124 inhibitor injected larvae may be due to expansion of the *Foxq2* expression domain (Fig. 6C), as observed in a different species of sea urchin that increased *Foxq2* expression domain leads to a decreased level of serotonin^30^. Thus, decreased serotonin may contribute to decreased gut contractions and swimming defects observed in miR-124 inhibitor injected embryos (Figs. 6A, 6C).

In addition, regulation of larval swimming is in part due to miR-124’s direct suppression of *NeuroD1*, since embryos injected with miR-124 inhibitor, *NeuroD1* TP, or exogenous *NeuroD1* transcripts, all lead to decreased swimming velocity (Figs 5A, 7C, S5B). The swimming defects is not likely attributed to structural defects of the cilia, as tubulin appears to be normal (Figs. 5C, 7E). Interestingly, we observed an increase in tubulin in miR-124 inhibitor injected and *NeuroD1* TP injected larvae (Figs. 5C, 7E). This may be due to miR-124’s direct suppression of *NeuroD1*, since it has been observed previously that an increase in NeuroD1 resulted in increased neuronal specific tubulin protein, TujI^102^. However, the exact mechanism of how increased NeuroD1 leads to increased cilia and neuronal tubulin is not known. Together, these results indicate that miR-124’s direct suppression of *NeuroD1* impacts larval swimming.

We found that miR-124 inhibitor injected larvae had a significant decrease in SynB expressing neurons compared to the control (Fig. 6B). This can be a result of significant loss of *Onecut* expression (Fig. 5D). *Onecut* is important in establishing oral-aboral polarity and in forming the ciliary band that allows for proper neural differentiation and neurite patterning^33^. During neurogenesis, SynB expressing neurons will differentiate throughout the band of *Onecut*-expressing cells, a process that has been found to be regulated by the inhibition of Nodal and BMP2/4 signaling^33^. Decreased *Onecut* expression has been shown to result in decreased neuronal bundling and proper interconnecting axonal tracts in sea urchin larvae^30^. Thus, decreased *Onecut* expression in miR-124 inhibitor injected embryos is likely to contribute to the decreased amount of mature neuronal connections (Figs. 5D and 6B). We do not know how miR- 124 mediates *Onecut* expression.

To reveal the molecular mechanism of how miR-124 regulates neurogenesis, we systematically assessed the spatial and temporal expression of key factors in the neuronal GRN in miR-124 perturbed embryos. Results indicate that miR-124 inhibition resulted in a 2-fold decrease of *Wnt6* and *FGFA*’s expression compared to the control, indicating that miR-124 is likely to regulate neurogenesis at an early stage when Wnt6 and FGFA specify the neuroectoderm domain in the blastula stage (Fig. 6D). Since neither Wnt6 or FGFA contains potential canonical miR-124 binding site, miR-124 is likely to regulate an activator of *Wnt6* and *FGFA* (Fig. 6D). Consistent with this hypothesis, Wnt6 is known to restrict *Foxq2* domain^7^, so a decrease in *Wnt6* transcripts in miR-124 inhibitor injected embryos may lead to an expansion of *Foxq2* expression domain (Fig. 6C, 6D).

Our results indicate that miR-124’s regulation of neurogenesis is broad, spanning from specification, differentiation, and maturation of neurons, whereas miR-124’s regulation of *NeuroD1* is specifically during the transition of neural differentiation to maturation. For the most part, *NeuroD1* TP phenocopies miR-124 inhibitor induced defects. However, a difference between miR-124 inhibitor and *NeuroD1* TP injected embryos is their *Foxq2* expression. While miR-124 inhibitor injected embryos have expanded expression domain of *Foxq2*, *NeuroD1* TP injected embryos did not have a change in the expression domain of *Foxq2* (Figs. 6C, 8C). In addition, miR-124 inhibitor injected gastrulae had a significant increase in the number of *SoxC* and *Brn1/2/4* expressing cells, indicating that miR-124 regulates neuronal specification to differentiation transition (Fig. 6E). miR-124’s impact on increased *SoxC* and *Brn1/2/4* expressing cells is likely due to its regulation of an additional unknown factor and not due to its regulation of *NeuroD1*, since *NeuroD1* TP injected blastulae and gastrulae did not have significant changes in the number of *SoxC* expressing cells and a net change of one additional *Brn1/2/4* expressing cell compared to the control (Fig. 8D, 8E). This is consistent with *NeuroD1*’s expression which peaks at the gastrula stage (Fig. 1B). Thus, from the miR-124 inhibition, *NeuroD1* knockdown and *NeuroD1* TP results, NeuroD1 is likely to regulate neuronal factors mainly at the late gastrula to larval stages during neuronal differentiation and maturation stages.

One additional discrepancy between the miR-124 inhibitor and *NeuroD1* TP injected embryos is that the number of *Elav* expressing cells in the miR-124 inhibitor injected larvae was significantly decreased compared to its control, while the *NeuroD1* TP injected embryos had increased *Elav* positive cells compared to its control larvae (Figs. 6F, 8F). The increase in *Elav* positive cells in the *NeuroD1* TP embryos is consistent with decreased *Elav* expression in NeuroD1 MASO, indicating that NeuroD1 positively regulates *Elav* (Figs. 1D, 8F). miR-124 has been observed in *Xenopus laevis* embryos to inhibit *NeuroD1* in the forebrain and optic vesicle, which resulted in an increased number of cells undergoing mitosis^46^. In addition, NeuroD1 promotes formation of neuron-like progenitor cells and has the ability to convert epithelial cells to the neural fate^50, 53^.^50, 51, 53, 71^. Thus, the increased number of *Elav* expressing cells in *NeuroD1* TP injected larvae may be due to NeuroD1’s function in enhancing proliferation and/or promoting formation of neuron-like cells that may not be functional (Fig. 8F). This is consistent with our results that despite an increase in *Elav* expressing cells, *NeuroD1* TP injected larvae have an overall loss of SynB positive neurons that are mature and functional (Fig. 8B).

To further reveal the mechanism that explains why *NeuroD1* TP injected embryos have increased *Elav* expressing cells and concomitant decrease of SynB-positive neurons, we hypothesize that miR-124’s inhibition of *Notch* may be critical for preventing excessive Delta/Notch mediated mitosis and apoptosis^15, 22, 102^. To test this, we coinjected *NeuroD1* TP and *Notch* TP into newly fertilized eggs and observed that these larvae have similar number of *Elav* positive cells as miR-124 inhibitor injected larvae (Fig. 8G). These results indicate that miR-124’s suppression of *Notch* is in part responsible for the decrease in *Elav* expressing cells in miR-124 inhibitor injected embryos (Fig. 8G). Blocking miR-124’s suppression of *Notch* may promote progenitor neurons to undergo lateral inhibition during the last mitotic division and induce apoptosis in non-neuronal cells^15, 22^. These data suggest that miR-124 regulates both *Notch* and *NeuroD1* to mediate proper neurogenesis.

Overall, we have integrated NeuroD1 into the neuronal network and determined that miR-124 regulates specification, differentiation, maturation, and function of neuronal development in part by mediating *Notch* and *NeuroD1* (Fig. 9). Specifically, miR-124 regulates an unidentified factor that activates *Wnt6, FGFA*, and *Foxq2* during neuronal specification. miR-124 also regulates another unknown factor that regulates *SoxC* and *Brn1/2/4* during neuronal differentiation in the gastrula stage. miR-124 represses *Notch* to ensure that the last mitotic division occurs properly. In the late gastrula to larval stages, miR-124 suppresses *NeuroD1* to mediate the transition between differentiation and maturation. miR-124 may suppress *NeuroD1* at the larval stage to prevent NeuroD1 from converting too many cells into neurons and allowing for the already committed neuronal cells to mature. Using the sea urchin embryo, we are able to systematically integrate miR-124’s post transcriptional regulation of the neuronal GRN and reveal miR-124’s mechanism of regulation. Overall, we identify miR-124 to have a prolific regulatory role throughout neurogenesis. Since miR-124, NeuroD1, and

**Figure 9.**
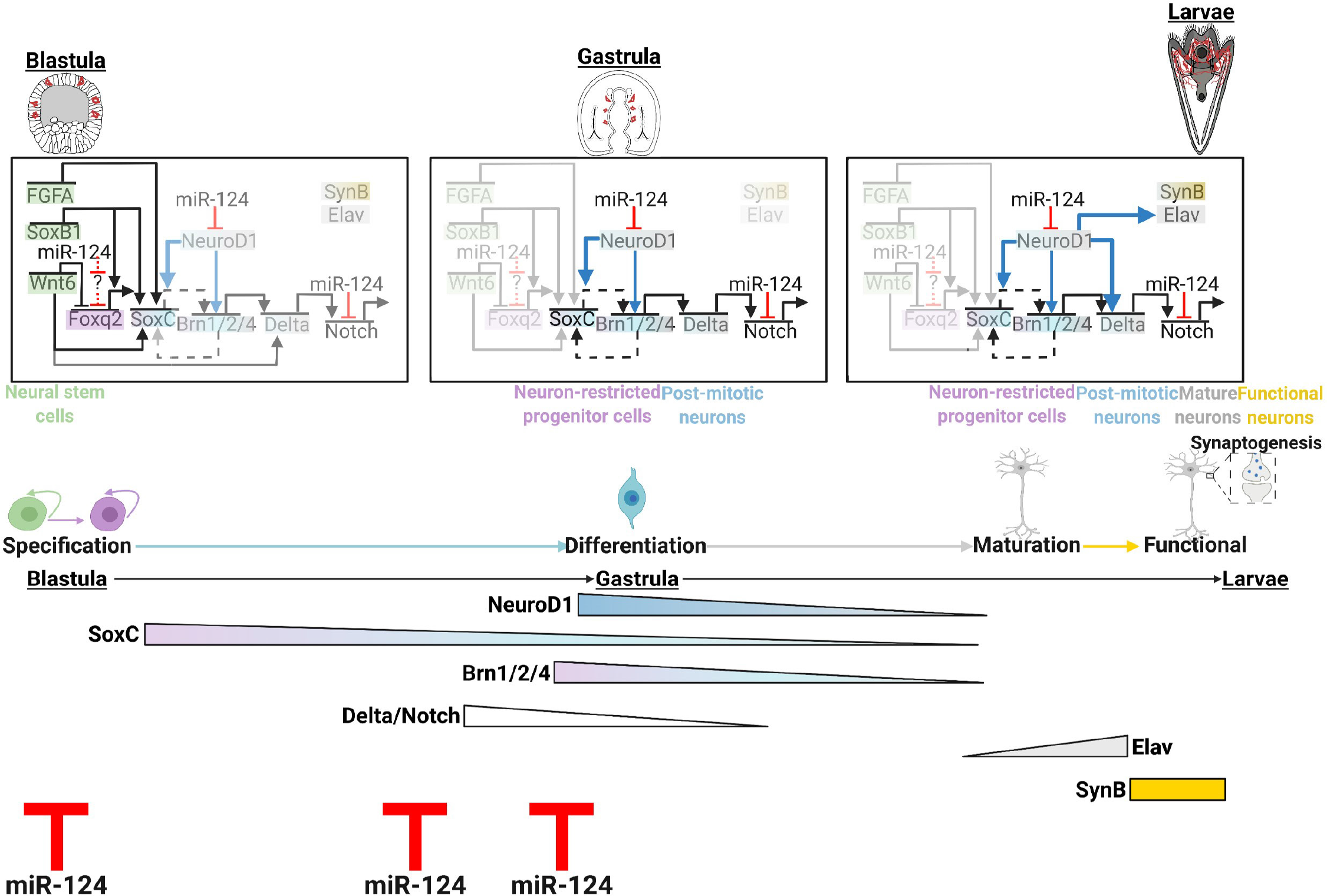
Working model of post-transcriptional control mediated by miR-124 in modulating neurogenesis. miR-124 regulates an unknown factor early in development to cause expression changes in *Wnt6*, *FGFA*, and *Foxq2* to mediate neuronal specification in the blastula stage. During the gastrula stage, miR-124 is likely to regulate an unidentified factor that regulates SoxC and Brn1/2/4 to mediate neuronal differentiation. miR-124 also directly inhibits *Notch* to allow neuronal progenitor cells to undergo the last mitotic division to become differentiated. *NeuroD1* expression peaks during the gastrula stage to perform its function mostly during the larval stage. During the late gastrula stage, NeuroD1 activates *SoxC, Brn1/2/4,* and inhibits *Delta*. Later during the larval stage, NeuroD1 activates *SoxC*, *Brn1/2/4, Delta* and *Elav*. miR-124 suppresses *NeuroD1* to modulate neuronal differentiation and maturation during the late gastrula and larval stages. miR-124 temporally regulates *Notch* and *NeuroD1* to facilitate neuronal differentiation and maturation. Our model is based on our observations. The Delta/Notch signaling is positioned according to prior published work^15, 22^. Made with Biorender.com

Notch are evolutionarily conserved, we believe that these results are applicable to our understanding of neurogenesis in other animals.

## Materials and methods

### Animals

Adult *Strongylocentrotus purpuratus* were collected from California coast (Point Loma Marine Company or Marinus Scientific, LLC.). All animals and cultures were cultured in 15°C incubator.

### Quantitative, real time polymerase chain reaction (qPCR)

To assess the relative quantity of *NeuroD1* transcripts throughout development, we collected 200 embryos at different development stages and extracted total RNA using the Micro RNeasy Kit (Qiagen, Germantown, MD). For NeuroD1 MASO, miR-124 inhibitor, and *NeuroD1* TP injected embryos, 100 embryos were collected and RNA extracted. cDNA was synthesized using the iScript cDNA Synthesis Kit (Bio-Rad, Philadelphia, PA). qPCR was performed using two embryo-equivalents for each reaction using Power SYBR Green PCR Master Mix (Invitrogen, Waltham, MA) in the Quantstudio6 Real-time PCR machine (Applied Biosystems, Waltham, MA) as previously described^96^. Primers were designed using the Primer3 program^103^ (Table 1).

**Table 1.**
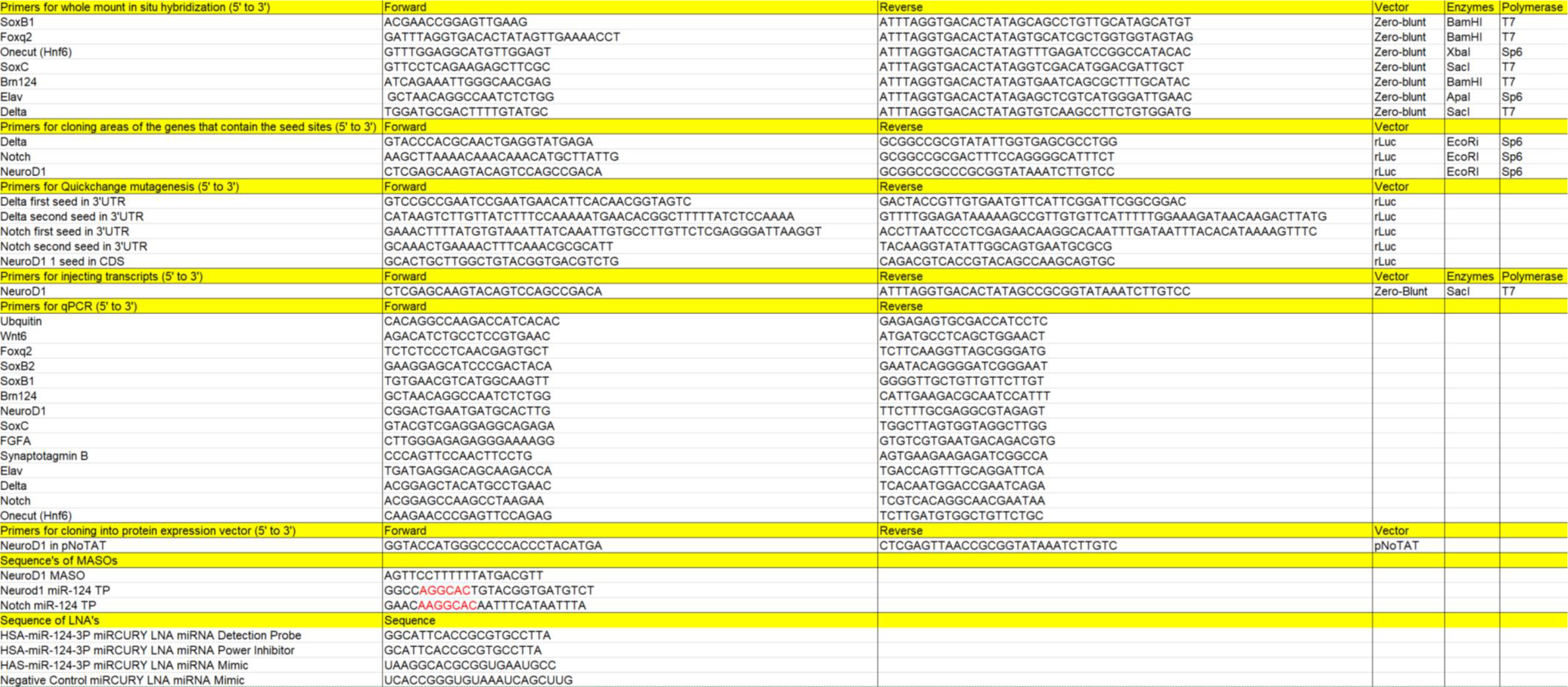

### Microinjections

Microinjections were performed as previously described with modifications ^66, 96, 104^. *Hsa*-miR-124-3p Locked Nucleic Acid (LNA) power inhibitor and *has*-miR-124- 3p miRCURY LNA miRNA mimic (Qiagen, Germantown, MD) were resuspended with RNase-free water to 100 μM. All sequences are listed in Table 1. Embryos were injected with the different concentrations (10 μM, 15 μM, and 20 μM) of miR-124 LNA inhibitor and collected at gastrula stage to phenotype for developmental defects. Based on the dose-response results, we used 15 µM of the *Hsa*-miR-124-3p LNA power inhibitor for subsequent experiments. The *Hsa*-miR-124-3p miRCURY LNA miRNA mimic (Qiagen, Germantown, MD) was used at 15µM with miR-124 inhibitor. To block miR-124’s binding and regulation of *NeuroD1*, we designed *NeuroD1* TP MASO against the miR-124 site within NeuroD1’s (Coding sequence) CDS (GeneTools, LLC, Philomath, OR)^97^. *NeuroD1* TP and control MASO (human *beta globin*) was resuspended to a 5 mM stock solution with RNAse-free water and diluted to 15μM, 30μM, and 300μM to perform microinjections. Zygotes were injected at the different concentrations and were phenotyped for defects. Translational blocking MASO against NeuroD1 (GeneTools, LLC, Philomath, OR) was resuspended to a 5 mM stock solution with RNAse-free water and diluted to 6 μM, 30 μM, and 150 μM to perform loss-of-function studies. Based on the dose-response results, we used 30 µM of the NeuroD1 MASO, where we observed ∼50% of injected blastulae survived, for subsequent experiments.

Injection solutions contained 20% sterile glycerol, 2 mg/mL 10,000 MW Texas Red or FITC lysine charged dextran (ThermoFisher Scientific, Waltham, MA) and various concentrations of miR-124 inhibitor, miR-124 mimic, NeuroD1 translational blocking MASO, or *NeuroD1* TP MASO. Injections were performed using the Pneumatic PicoPump with vacuum (World Precision Instruments; Sarasota, FL)^96, 105^. A vertical needle puller PL-10 (Narishige, Tokyo, Japan) was used to pull the injection needles (1 mm glass capillaries with filaments) (Narishige Tokyo, Japan).

### RNA in situ hybridization

The steps performed for fluorescence RNA *in situ* hybridization (FISH) are described previously with modifications^106^. All sequences are listed in Table 1. The Dre- miR-124-3p miRCURY LNA detection probe Qiagen, Germantown, MD) was used to visualize sea urchin miR-124 (at 0.5 ng/µl in hybridization buffer and incubated at 50°C for five days. The scrambled-miR miRCURY LNA detection probe was used as a negative control at the same concentration as miR-124 probe.

To generate RNA probes against protein coding transcripts, we used PCR to amplify *Onecut (Hnf6), SoxB1, SoxC, Delta, Foxq2, NeuroD1, Brn1/2/4,* and *Elav* from sea urchin egg and embryonic cDNA of 24 hpf, 48 hpf, 72 hpf. PCR primers and enzymes used to linearize and make antisense probes are listed in Table 1. Other probes including *FoxA, Krl,* and *Vasa* were previously cloned^107, 108^. 0.5 µg probe/mL was used to detect native transcript in embryos, according to previous protocols ^105^. The embryos are incubated with the 1:1,000 anti-digoxigenin-POD antibody overnight at 4°C and amplified with Tyramide Amplification working solution and exposed with fluorescein (1:150 dilution of TSA stock with 1x Plus Amplification Diluent-fluorescein) (Akoya Biosciences, Marlborough, MA). All FISH embryos were mounted using DAPI in MOPs buffer and NucBlue (ThermoFisher Scientific, Waltham, MA). Images were taken using the Zeiss LSM 880 scanning confocal microscope (Carl Zeiss Incorporation, Thorwood, NY). The maximum intensity projections of Z-stack of images were acquired with Zen software and exported into Adobe Photoshop (Adobe, San Jose, CA) for further image processing.

The steps performed for colorimetric whole mount *in situ* hybridization (WMISH) are as previously described^105, 109^. Embryos were incubated 1:1500 anti-digoxigenin-Alkaline phosphatase antibody overnight at room temperature. Embryos were imaged using the Observer Z.1 microscope (Carl Zeiss Incorporation, Thorwood, NY). The Z-stack slice at the equatorial plane was taken and exported into Adobe Photoshop (Adobe, San Jose, CA) for further image processing.

### Immunolabeling procedures

To access miR-124 inhibitor induced phenotypes, we used antibodies against various cell types. We used Endo1 to detect mid- and hindgut^73^, E7 (Developmental Studies Hybridoma Bank (DSHB), Lot#2/13/20-54µg/mL^110^) to detect tubulin in cilia, 1E11 (DSHB, Lot #3/26/14-30 µg/mL) and SynB (from Gary Wessel, Brown University) to detect SynB-expressing neurons^26, 32, 87^, serotonin (MilliporeSigma, St. Louis, MO cat# S5545) ^80, 111^, NeuroD1 (ABclonal, Woodburn, MA cat# A1147). Embryos were fixed in 4% paraformaldehyde (PFA) (20% stock; EMS, Hatfield, PA) in artificial sea water overnight at 4°C. Three 15-minute Phosphate Buffered Saline-Tween-20 0.05% (PBST) (10X PBS; Bio-Rad, Hercules, CA) washes were performed. Embryos were blocked with 4% sheep serum (MilliporeSigma, St. Louis, MO) for 1 hour at room temperature.

For 1E11 and NeuoD1, the embryos were fixed in 4% PFA for 10 minutes and post fixed with 100% acetone for 1 minute and washed with PBST containing 0.1% Triton X- 100 (Thermo Fisher Scientific, Waltham, MA) and incubated for two nights at 4°C in blocking buffer (10% Bovine serum albumin (MilliporeSigma, St. Louis, MO) in PBST 0.1% Triton X-100). For SynB antibody, we fixed embryos 3.7% formaldehyde in filtered natural water (FSW) for 20 minutes at room temperature and 1 minute post fix with ice cold methanol and washed with PBST-0.1% Triton X-100 and incubated overnight.

Primary antibody incubation was performed with Endo1, 1E11, SynB, serotonin, and NeuroD1, at 1:50, 1:2, 1:200, 1:500, and 1:250, respectively. Embryos were washed three times 15 minutes with PBST followed by incubation with secondary antibodies goat anti-mouse (for Endo1 and 1E11) and goat anti-rabbit (SynB, Serotonin, NeuroD1) Alexa 488 or Alexa 647 at 1:300 for 1 hour at room temperature (Thermo Fisher Scientific, Waltham, MA). For tubulin immunolabeling, control embryos were injected with a non-fixable FITC and the miR-124 inhibitor injected embryos were injected with fixable Texas Red. Control and miR-124 inhibitor injected embryos were immunolabeled with tubulin in both separate or the same well to make sure differences observed are not due to potential technical differences.

The Phalloidin conjugated to Alexa 488 (ThermoFisher Scientific, Waltham, MA) was resuspended in 100% methanol to make 200 U/ml stocks and then lyophilized and re-suspended in 1XPBS-0.1% Titon-X-100 to make a final concentration of 10 U/ml, which was added to the embryos. Embryos were fixed in 4% PFA in 1XPBS for 5 min on ice and then placed at room temperature for 15 minutes. They were post-fixed in 100% acetone for 10 min on ice and washed with PBST-0.1% Triton-X-100, followed by incubation with Phalloidin for 1 hour at room temperature and three washes with PBST and then 1XPBS. All immunolabeled embryos were imaged using a Zeiss LSM 880 scanning confocal microscope (Carl Zeiss Incorporation, Thorwood, NY). All immunolabeled embryos were mounted using DAPI in PBST buffer (NucBlue; ThermoFisher Scientific, Waltham, MA). The maximum intensity projections of Z-stack of images were acquired with Zen software (Carl Zeiss Incorporation, Thorwood, NY) and exported into Adobe Photoshop (Adobe, San Jose, CA) for further processing.

### Quantification

To measure the levels of miR-124, 1E11, serotonin, tubulin and NeuroD1 (protein and RNA) in each embryo, we took maximum intensity projections and exported them into ImageJ^110^. The serotonin and 1E11 containing region in the ciliary band was measured and the background was subtracted. For miR-124 levels, that average fluorescent intensity was calculated by measuring the area of the embryo of interest and subtracting the average fluorescent background.

To measure gut contraction, we mounted the embryos with protamine sulfate coverslips in FSW. The sides of the coverslip with sealed with melted petroleum jelly to allow airflow. Each embryo was recorded for four minutes, and the number of contractions were counted using Observer Z.1 microscope (Carl Zeiss Incorporation, Thorwood, NY) 40X lens. The gut contraction was determined by the foregut opening into the midgut as a full contraction.

To track swimming movement, we used the manual tracking plugin in ImageJ^110^. We set the time interval at 60 seconds and the x/y calibration at 0.645 µm. Each movie was imaged for 60 seconds and only embryos that stayed in the field of view were analyzed. We track the leading edge of the larvae which was the top of the mouth and followed through the entirety of the movie. To compose the movies, we used four frames per second, consisting of a total of a fifteen second video.

To assess the beating of the cilia, polybead dyed blue 1µm microspheres (Polyscience Inc, Warrington, PA) were used. Embryos were injected with either FITC or Texas Red dextrans and mounted on the same coverslip to limit variability of beads between control and perturbed embryos. The polybeads were used at 1:500 in sea water. Prior to its use, the beads were sonicated in a water bath for 20 minutes. We mounted the control and experimentally treated embryos with the diluted polybeads and image them using Observer Z.1 microscope (Carl Zeiss Incorporation, Thorwood, NY) for 2 minutes. To quantify the ciliary beating flow videos, we drew a rectangle size of 15 µm x 60 µm that was 15 µm away from the mouth of the embryo. The beads were counted as they entered the imaging area, and the number of beads was normalized to the control. Before normalization, the average number of beads for the control and experimentally treated embryos was subtracted by the average number of beads from the deciliated larvae. To determine that the flow of the beads was due to the actual cilia beating and not due to some other external factor, we took a set of physiological embryos and deciliated them with 2x sea water for five minutes and washed off with normal sea water and mounted with beads. The number of beads was counted as described in methods. Then the average number of beads collected from the deciliated larvae was subtracted from the average number of beads from the control and experimental embryos.

### Cloning of constructs for luciferase assays

The CDS of *NeuroD1* and 3’UTRs of *Delta* and *Notch* was cloned using sea urchin cDNA into Zeroblunt vector (ThermoFisher Scientific, Waltham, MA) (Table 1). Plasmids containing potential cloned DNA inserts were subjected to DNA sequencing (Genewiz Services, South Plainfield, NJ). *NeuroD1* CDS, *Delta* 3’UTR, and *Notch* 3’UTR was subcloned downstream of the Renilla luciferase (*RLuc)* as described previously^105^. miR-124 seed sequence was deleted from the *NeuroD1* CDS, by using the QuikChange Lightning Kit (Agilent Technologies, San Jose, CA). The seeds within the 3’UTR of *Delta* and *Notch* were mutagenized at the third and fifth binding site^112^.

The sequence of the mutagenesis primers used are listed in Table 1. Clones were sequenced to check for the deleted or mutated miR-124 binding site (Genewiz Services, South Plainfield, NJ). *NeuroD1 RLuc* reporter constructs primers and restriction enzymes and RNA polymerases used are listed in Table S2. Firefly construct (*FF)* was linearized using *SpeI* and *in vitro* transcribed with SP6 RNA polymerase^105^. Transcripts were purified using the RNA Nucleospin Clean up kit (Macherey-Nagel, Bethlehem, PA). *FF* and reporter *RLuc* constructs were co-injected at 50 ng/µL. 50 embryos at the mesenchyme blastula stage (24 hpf) were collected in 25 µL of 1X Promega passive lysis buffer and vortexed at room temperature. Dual luciferase assays were performed using the Promega™ Dual-Luciferase™ Reporter (DLR™) Assay Systems with the Promega™ GloMax™ 20/20 Luminometry System (Promega, Madision, WI). The rest of the assay was performed as previously described^105^.

### Preadsorption assay of NeuroD1

To test if the NeuroD1 antibody developed against the human antigen would cross react with the sea urchin NeuroD1, we first cloned the sea urchin NeuroD1 into an expression vector pNOTAT^113^. Plasmids were sequenced (University of Delaware DNA Sequencing and Genotyping Center, Newark, DE and Genewiz, South Plainfield, NJ).

NeuroD1-pNOTAT was transformed into C41 competent cells (Lucigen, Middleton, WI) and induced with IPTG at 0.1mM for protein expression. Negative control and NeuroD1 lysates with or without IPTG induction were run on an 12% SDS-PAGE gel. The protein gel was transferred to the PVDF membrane (Immun-Blot PVDF Membranes for Protein Blotting, BioRad, Philadelphia, PA) using a wet transfer system (Wet Tank Transfer Systems, BioRad, Philadelphia, PA). The blot was blocked in 3% Bovine serum albumin (Research Products International, Mount Prospect, IL) in TBST (50mM Tris, pH 7.5, NaCl 184mM, Tween-20 0.05%) for 1 hour at room temperature. Then NeuroD1 antibody was incubated overnight at 4°C at 1:1,000 concentration. Blot was exposed with SuperSignal West Pico Chemiluminescent Substrate (ThermoFisher Scientific, Waltham, MA) and imaged with Biorad Chemidoc Gel Imager (BioRad, Philadelphia, PA).

As one test of specificity of the antibodies, a preadsorption assay was performed by incubating the NeuroD1 antibodies with the bacterial lysate containing the sea urchin NeuroD1 protein. First, 50µL of prepared bacterial lysate was incubated with the PVDF membranes (0.5 cm x 0.5 cm) in the Eppendorf tubes and rocked for two nights at 4°C. Then the NeuroD1 antibody was added at 1:250 in blocking buffer and incubated with the PVDF containing negative control or the sea urchin NeuroD1 for two nights at 4°C in blocking in PBST- 0.1 %Triton in 4% sheep serum. NeuroD1 antibody preadsorbed with bacterial lysate with or without sea urchin NeuroD1 was used in immunolabeling experiment with fixed sea urchin larvae. Embryos were washed with PBST and exposed with goat anti-rabbit 488 in blocking buffer (4% sheep serum) for one hour at room temperature and counterstained with DAPI. Images were acquired with Zeiss 880 confocal microscope.

### Preparation of RNA transcripts for injections

NeuroD1 CDS was *in vitro* transcribed with Sp6 (mMessage, ThermoFisher Scientific, Waltham, MA). Transcripts were purified using the RNA Nucleospin Clean up kit (Macherey-Nagel, Bethlehem, PA). The transcribed *NeuroD1* was injected at 1, 2, and 3 µg/µL with cytoplasmic *mCherry* RNA as control^114^ at 3, 2, and 1 µg/µL, respectively. The control injection solution contained 4 µg/µL of *mCherry*, which allowed us to detect potential RNA degradation. The RNA was passed through a spin column (MilliporeSigma, St. Louis, MO) prior to injection. *NeuroD1* transcript and *mCherry* control were injected at 1µg/µL or subsequent experiments, based off the dose response experiment.

## Supporting information

Control mimic injected larvae were imaged live for 60 seconds and tracked manually with ImageJ plugin to obtain the velocity.

miR-124 inhibitor injected larvae were imaged live for 60 seconds and tracked manually with ImageJ plugin to obtain the velocity.

miR-124 inhibitor and miR-124 mimic were coinjected and imaged live for 60 seconds and tracked manually with ImageJ plugin to obtain the velocity.

Control injected embryos were imaged lived for cilia beating for 120 seconds with polybeads. Each video is composed of 4 frames per second.

Embryos were deciliated with 2X sea water and imaged lived for 120 seconds with polybeads. Each video is composed of 4 frames per second.

miR-124 inhibitor injected larvae were imaged lived for cilia beating for 120 seconds with polybeads. Each video is composed of 4 frames per second.

Control TP injected embryos were imaged for four-minute period and counted gut contractions in foregut. Each video is composed of 4 frames per second.

NeuroD1 TP injected embryos were imaged for four-minute period and counted gut contractions in foregut. Each video is composed of 4 frames per second.

Control TP injected embryos were imaged live for 60 seconds and tracked manually with ImageJ plugin to obtain the velocity.

NeuroD1 TP injected embryos were imaged live for 60 seconds and tracked manually with ImageJ plugin to obtain the velocity.

Control TP injected embryos were imaged live for cilia beating for 120 seconds with polybeads.

NeuroD1 TP injected larvae were imaged live for cilia beating for 120 seconds with polybeads.

mCherry injected embryos were imaged for four-minute period and counted gut contractions in foregut. Each video is composed of 4 frames per second.

NeuroD1 transcript injected embryos were imaged for four-minute period and counted gut contractions in foregut. Each video is composed of 4 frames per

NeuroD1 transcripts (NeuroD1 OE) injected embryos were imaged live for 60 seconds and tracked manually with ImageJ plugin to obtain the velocity.

mCherry control injected embryos were imaged live for 60 seconds and tracked manually with ImageJ plugin to obtain the velocity.

Control injected embryos were imaged for four-minute period and counted gut contractions in foregut. Each video is composed of 4 frames per second.

miR-124 inhibitor injected embryos were imaged for four-minute period and counted gut contractions in foregut.

## Acknowledgments

We thank Dr. Gary Wessel (Brown University) for the Endo1 antibody and SynB antibody. We want to thank the anonymous reviewers for their time and valuable feedback.

## Competing interests

The authors declare no competing or financial interests.

## Author contributions

K.D.K. and J.L.S. conceived the experiments. K.D.K. performed and analyzed all the experiments. K.D.K and J.L.S. wrote the paper.

## Funding

This work is funded by National Science Foundation (IOS 1553338 to J.L.S.), NIH P20GM103653, NIH NIGMS P20GM103446, and Sigma Xi Grants-in-Aid of Research (G2018031596227966 to K.D.K.).

---

**Figure S1.**
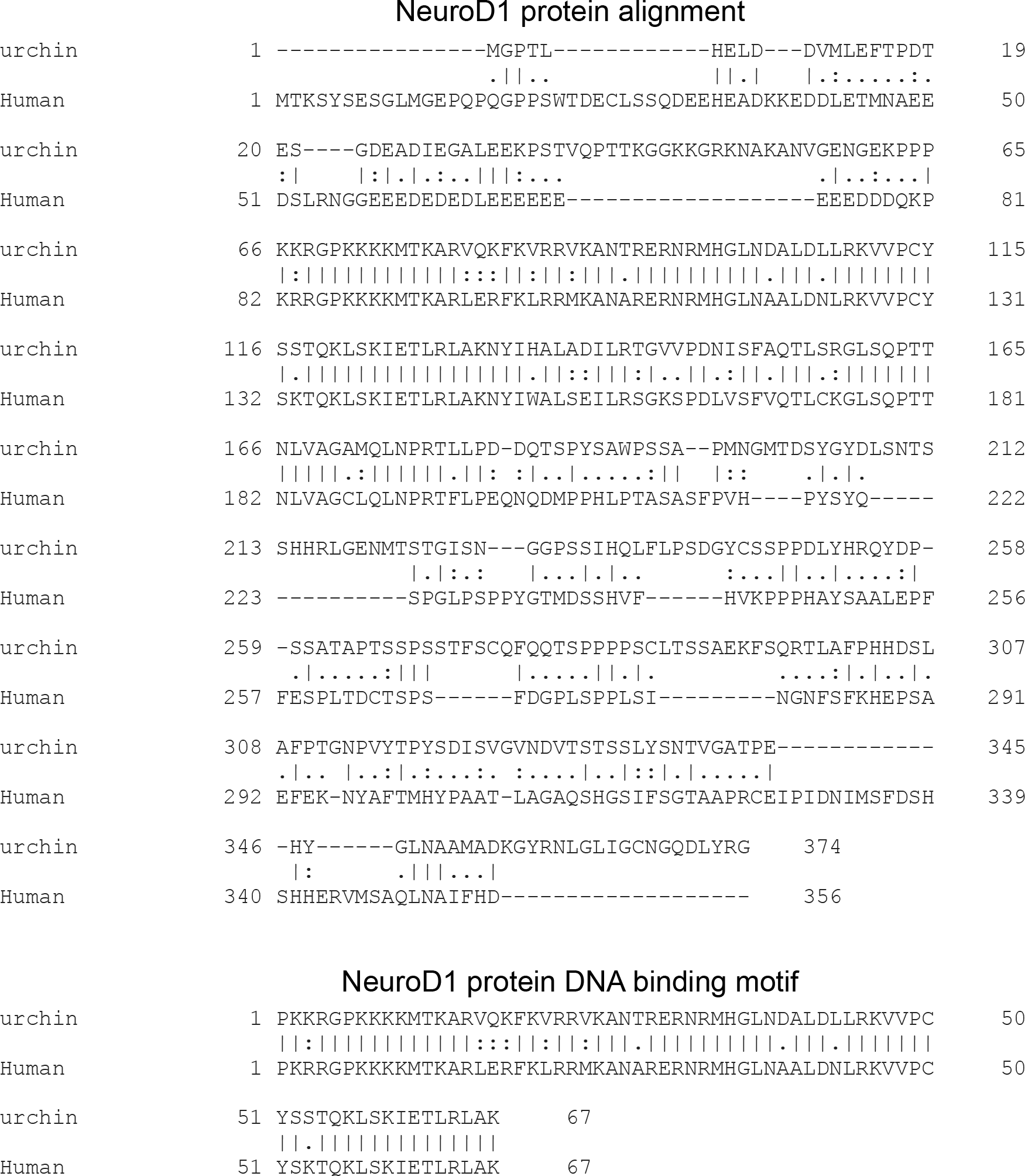
NeuroD1 is conserved between human and sea urchin. Using Clustal Omega, we aligned the protein sequences from the sea urchin and human NeuroD1. Using National Center for Biotechnology Information (NCBI) protein alignment, we determined that there was a 33.7% identity between human and sea urchin NeuroD1 protein sequence and an 85.1% identify within the DNA binding domain.

**Figure S2.**
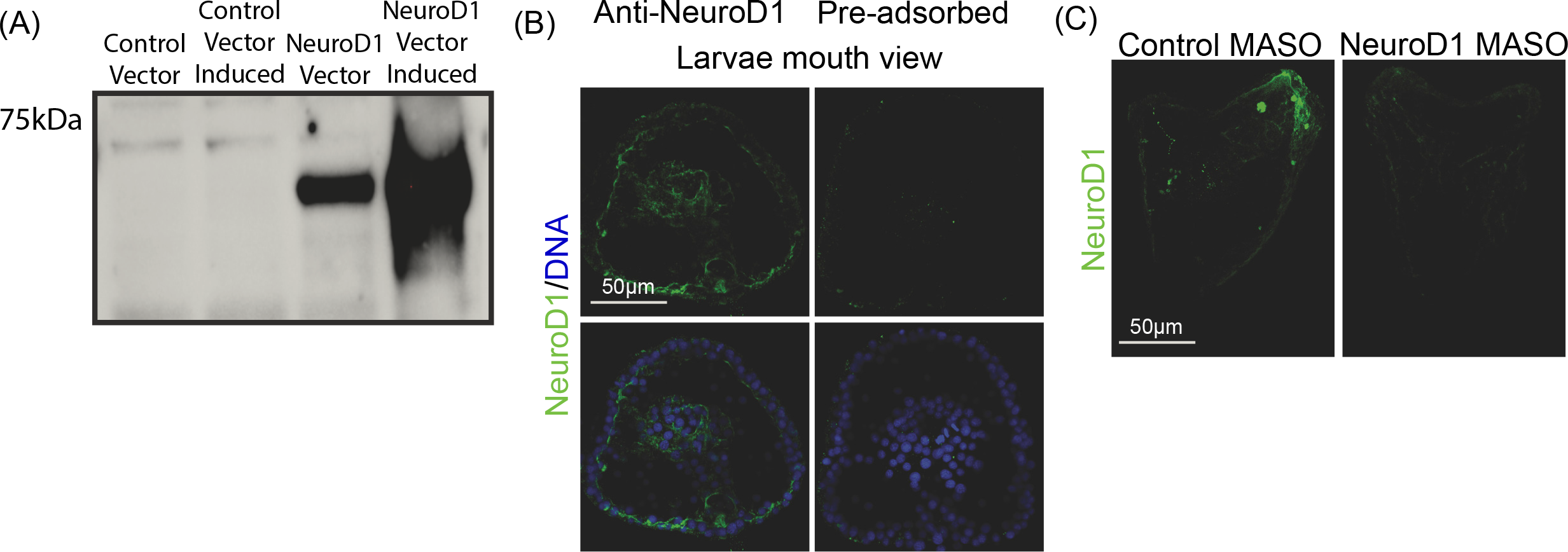
Human NeuroD1 antibody cross reacts with the sea urchin NeuroD1. (A) Western blot of bacterial lysates with vector alone or expression plasmid containing sea urchin NeuroD1 induced with or without IPTG. (B) Testing of the specificity of the NeuroD1 antibody with preadsorption assay. Larvae were immunolabeled with pre-adsorbed NeuroD1 with or without sea urchin NeuroD1 protein (green) and counterstained with DAPI (blue). (C) Embryos injected with NeuroD1 MASO (30 µM) had significantly less NeuroD1 protein (green) compared to the control

**Figure S3.**
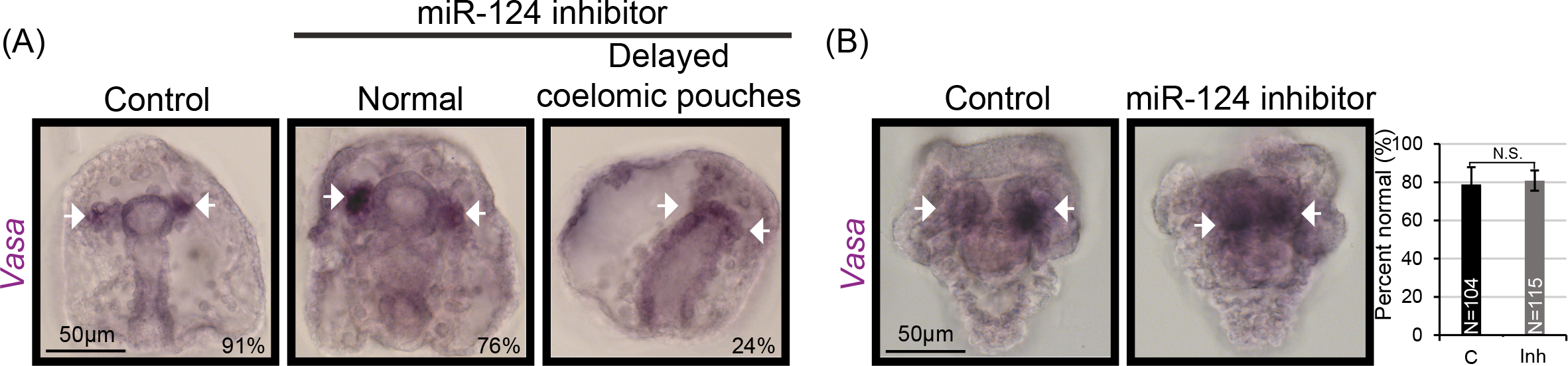
miR-124 inhibition results in delayed mesodermally derived coelomic pouch formation. (A) 24% of miR-124 inhibitor injected gastrulae did not form visible coelomic pouches but had *Vasa*-positive multipotent cells (purple stain). Control N=33 and miR-124 inhibitor injected embryos N=29. White arrows point to *Vasa*-positive multipotent cells. (B) miR-124 inhibitor injected larvae and control both had visible *Vasa*- positive coelomic pouches. 3 biological replicates. SEM is graphed. Cochran-Mantel-Haenszel statistical test was used. Black arrows point to the *Vasa*-positive multipotent cells.

**Figure S4.**
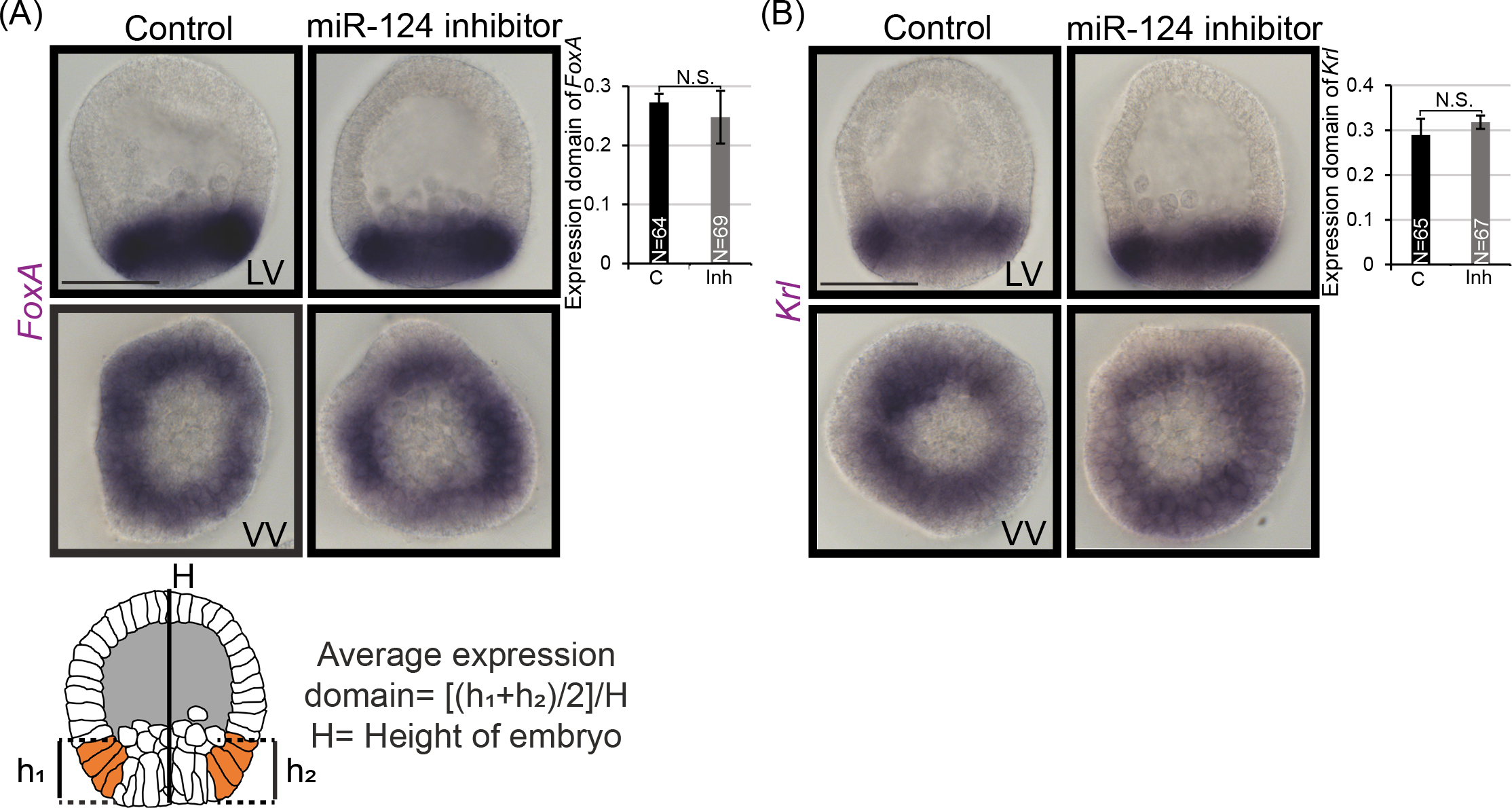
miR-124 inhibition results in no change in endoderm specific genes. (A) *FoxA* expression was not altered in miR-124 inhibitor injected blastulae compared to the control (purple stain). Schematic of the measurements used to analyze the expression domain is depicted. 3 biological replicates. (B) *Krl* expression was not altered in miR-124 inhibitor injected blastulae compared to control. 3 biological replicates. N= total number of embryos. LV= lateral view. VV=vegetal view. SEM is graphed. Student t-test was used.

**Figure S5.**
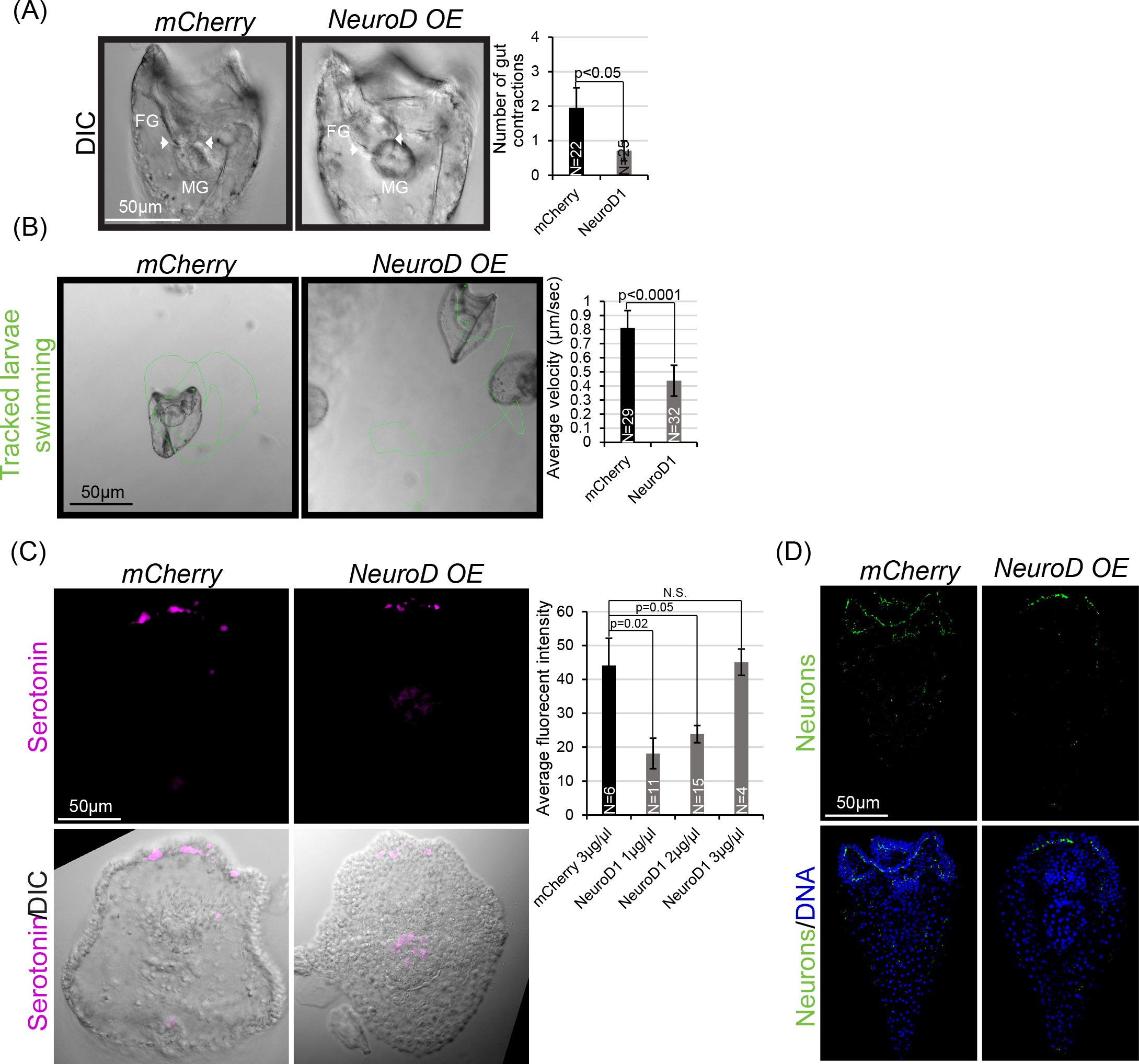
NeuroD1 overexpression results in decreased gut contractions and swimming velocity. (A) Injection of *NeuroD1* transcripts into the sea urchin zygotes resulted in a significant decrease in gut contractions compared to the control. Still frame images of embryos during gut contractions are depicted. (B) NeuroD1 overexpression (OE) results in a significant decrease in swimming velocity compared to the control. (C) Oral view of serotonin-containing neurons (magenta) with close-up view (shown by the inset delineated by the white dashed lines). Larvae were immunolabeled with the serotonin antibody (magenta) and counterstained with DAPI (blue). 2 biological replicates. (D) NeuroD1 OE larvae were immunolabeled with sea urchin neuronal antibody 1E11 that detects SynB expressing neurons (green) and counterstained with DAPI (blue). NeuroD1 OE larvae exhibited decreased SynB positive neurons compared to the control. 3 biological replicates. Maximum intensity projection of Z-stack confocal images is presented.

**Video 1A.** Control injected embryos were imaged for four-minute period and counted gut contractions in foregut. Each video is composed of 4 frames per second.

**Video 1B.** miR-124 inhibitor injected embryos were imaged for four-minute period and counted gut contractions in foregut.

**Video 2A.** Control mimic injected larvae were imaged live for 60 seconds and tracked manually with ImageJ plugin to obtain the velocity.

**Video 2B.** miR-124 inhibitor injected larvae were imaged live for 60 seconds and tracked manually with ImageJ plugin to obtain the velocity.

**Video 2C.** miR-124 inhibitor and miR-124 mimic were coinjected and imaged live for 60 seconds and tracked manually with ImageJ plugin to obtain the velocity.

**Video 3A.** Control injected embryos were imaged lived for cilia beating for 120 seconds with polybeads. Each video is composed of 4 frames per second.

**Video 3B.** Embryos were deciliated with 2X sea water and imaged lived for 120 seconds with polybeads. Each video is composed of 4 frames per second.

**Video 3C.** miR-124 inhibitor injected larvae were imaged lived for cilia beating for 120 seconds with polybeads. Each video is composed of 4 frames per second.

**Video 4A.** Control TP injected embryos were imaged for four-minute period and counted gut contractions in foregut. Each video is composed of 4 frames per second.

**Video 4B.** *NeuroD1* TP injected embryos were imaged for four-minute period and counted gut contractions in foregut. Each video is composed of 4 frames per second.

**Video 5A.** Control TP injected embryos were imaged live for 60 seconds and tracked manually with ImageJ plugin to obtain the velocity.

**Video 5B.** *NeuroD1* TP injected embryos were imaged live for 60 seconds and tracked manually with ImageJ plugin to obtain the velocity.

**Video 6A.** Control TP injected embryos were imaged live for cilia beating for 120 seconds with polybeads.

**Video 6B.** *NeuroD1* TP injected larvae were imaged live for cilia beating for 120 seconds with polybeads.

**Video 7A.** mCherry injected embryos were imaged for four-minute period and counted gut contractions in foregut. Each video is composed of 4 frames per second.

**Video 7B.** *NeuroD1* transcript injected embryos were imaged for four-minute period and counted gut contractions in foregut. Each video is composed of 4 frames per second.

**Video 8A.** *mCherry* control injected embryos were imaged live for 60 seconds and tracked manually with ImageJ plugin to obtain the velocity.

**Video 8B.** *NeuroD1* transcripts (NeuroD1 OE) injected embryos were imaged live for 60 seconds and tracked manually with ImageJ plugin to obtain the velocity.

## Notes

### Competing Interest Statement

The authors have declared no competing interest.

